# Nek family members regulate Rad54 during homologous recombination in developing mice

**DOI:** 10.64898/2025.12.19.695536

**Authors:** Isabel Freund, Jie Liu, Holly Thomas, Mareike Haarmann, Emilie Renaud, Florian Frohns, Jia-Xuan Chen, Wolf-Dietrich Heyer, Markus Löbrich

## Abstract

Homologous recombination (HR) represents an important pathway for repairing DNA double-strand breaks (DSBs) but HR factors, including RAD51, also serve to protect and restart stalled replication forks. RAD54 functions during DSB repair where it removes RAD51 from duplex DNA including heteroduplex DNA which is formed during D-loop formation. This allows subsequent DNA repair synthesis and completion of the HR process. We have previously suggested that RAD54’s activity is regulated by never-in-mitosis-gene A (NIMA) related kinase 1 (NEK1) in a cell cycle-specific manner to promote RAD51 removal and HR in G2 phase without interfering with RAD51’s fork stabilization role during S phase. Here, we establish that Nek1 regulates the phosphorylation of Rad54 at Ser572 (S572) to promote HR in vivo in adult mice and in vitro in fibroblasts derived from such mice. In contrast, embryonic mice and fibroblasts derived from them do not require Nek1 for HR. We further show that HR requires Rad54 phosphorylation at S572 both in embryonic and adult fibroblasts and that this is mediated in embryonic fibroblasts by Nek3 and Nek5 instead of Nek1. Thus, our work identifies a developmental change in the regulation of HR and uncovers two new factors involved in this process.

## Introduction

DNA damage constantly arises over the lifespan of an individual and can have important cellular consequences. If left unrepaired, it can lead to cell death and if incorrectly repaired, mutations can arise.^1^ This not only contributes to cancer development and aging but also compromises developmental processes.^2–4^ Among the various damages that arise endogenously or from exposure to exogenous agents, DNA double-strand breaks (DSBs) are particularly harmful since sequence information is lost on both strands.^5^ A network of cellular responses has evolved to counteract DNA damage which is called the DNA damage response (DDR) and includes repair pathways, processes of cell cycle checkpoint regulation and apoptosis.^6^ Defects in the DDR pathways lead to genomic instability and confer susceptibility to cancer and degenerative diseases.^7^

Homologous recombination (HR) is one of two main pathways for repairing DSBs.^8^ HR repairs the damaged DNA by using the sister chromatid as a template for repair and is hence limited to the S and G2 phases of the cell cycle.^9^ The process is initiated by DNA end-resection generating single-stranded DNA (ssDNA) overhangs that are subsequently loaded with the key HR factor RAD51 by BRCA2.^10,11^ The resulting RAD51 nucleoprotein filament mediates homology search and promotes the pairing of the ssDNA end with one of the strands of the sister chromatid in a process called D-loop formation.^12^ Another important HR factor, RAD54, is required for this process and removes RAD51 from the generated heteroduplex DNA during D-loop formation.^13^ This step is critical for the subsequent DNA repair synthesis and the completion of HR. HR can also repair daughter strand gaps that arise behind the replication fork using the sister chromatid as a template.^14,15^ Importantly, many HR factors have additional roles during DNA replication when replication forks encounter DNA damage and stall.^16^ RAD51 promotes the reversal of stalled forks to allow damage repair and replication restart.^17^ Arrested and reversed forks need to be protected from nucleolytic degradation, a function that might be supported by the ability of yeast and human RAD51 to bind double-stranded DNA (dsDNA).^18,19^ The precise mechanisms operating at stalled forks are not yet completely understood, and it is unclear if and how RAD51’s role at stalled forks is regulated differently to its role during HR. One possibility is that the process of HR at DSBs and daughter strand gaps, and the function of HR proteins at the replication fork, are separated during the cell cycle. RAD54 might exert a key function in this regulation since its activation and phosphorylation by NEK1 (never-in-mitosis-gene A (NIMA) related kinase 1) occurs specifically during the G2 phase, irrespective of where in the cell cycle the damage was induced. This G2-specific activation of RAD54 ensures that RAD51 is not removed from stalled forks.^20^

The mammalian NEK family of serine/threonine kinases comprises 11 genes, *nek1-11*, which have been characterized based on their homology to the nimA gene of the fungus *Aspergillus nidulans*.^21,22^

While the N-terminal kinase domains are highly conserved, the NEK family members strongly differ in the structural composition of their C-termini. While some members, including NEK1, 2, 9, 10 and 11, harbor coiled-coil domains for protein interaction as well as PEST sequences for proteasomal degradation, NEK3 only contains PEST sequences and NEK5 only coiled-coil domains. NEK5 is unique among all NEKs in containing a central DEAD-box helicase-like domain.^23^ Given their structural diversity and complexity, it is not surprising that the NEK family members play an important role in more than one cellular process. Although only a few direct substrates have been identified so far, all NEKs are involved to some extent in primary cilia formation, cell cycle regulation, microtubule/centrosome organization and/or mitosis.^24^ Importantly, there is growing evidence that NEK kinases regulate several DDR processes, including cell cycle checkpoint induction, DNA repair and apoptosis.^25^

NEK1 is arguably the best characterized NEK family member and its complete loss has been associated with the human disorder *short-rib polydactyly syndrome (SRPS) type Majewski* which is embryonically lethal and has only been observed in aborted fetuses.^26^ A homozygous single-base insertion in the Nek1 mouse gene which causes the formation of a premature stop codon leads to a severe pleiotropic phenotype that manifests around 14 days after birth.^27^ These so-called kat2j mice exhibit male sterility, female fertility reduction and premature lethality caused by numerous developmental disorders including hydrocephalus, dwarfism, facial dysmorphia, anemia and polycystic kidney disease (PKD).^28,29^

Here, using a mouse model system, we discover that the requirement of Nek1 for HR depends on the developmental stage, confirming and extending our previous work with HeLa tumor cells. We establish that Nek1 regulates the phosphorylation of Rad54 at Ser572 (S572) to promote HR *in vivo* in adult mice and *in vitro* in fibroblasts derived from these mice. In contrast, embryonic mice and fibroblasts derived from them do not require Nek1 for HR. We further show that HR requires Rad54 phosphorylation at S572 both in embryonic and adult fibroblasts and that this is mediated in embryonic fibroblasts by Nek3 and Nek5 instead of Nek1. Thus, our work identifies a developmental change in the regulation of HR and uncovers two new factors involved in this process.

## Results

### Nek1-KO mice show unresolved Rad51 foci late but not early during development

We previously characterized the role of NEK1 as a regulator of RAD54 during HR in HeLa tumor cells and showed that NEK1-deficient cells are defective in HR and have elevated levels of RAD51 foci.^20^ To validate this function of Nek1 *in vivo*, we established a protocol that allows the cell cycle-dependent analysis of DSB repair via HR in paraffin-embedded mouse tissues. A detailed description of this procedure is provided in the supplement (Fig. S1). Briefly, wild-type (WT), Nek1 knock-out (Nek1-KO), and Rad54-KO mice of different developmental stages were treated with the thymidine analog EdU by intraperitoneal injection to mark S-phase cells, exposed to 1 Gy of X-rays, and sacrificed 2 or 8 h post IR. We initially analyzed cells of the brain since we previously established immunofluorescence analysis in this model tissue.^30,31^ Brains were fixed, coated with paraffin, and tissue sections stained for cell cycle markers and Rad51 as a readout for Rad54 activity since RAD51 foci have previously been shown to persist in RAD54-deficient HeLa cells.^20^ We identified proliferative cell populations in the hippocampus of the postnatal brain (P21, Fig. 1A), in the cerebellum of the neonatal brain (P4, Fig. 1B), and in the lateral ventricles of the embryonic brain (E14.5, Fig. 1C) (see also Fig. S2). We then assessed Rad51 foci numbers and the cell cycle distribution of cells that were, based on their EdU staining pattern, irradiated in mid-S phase. All mice displayed comparable amounts of Rad51 foci 2 h post IR, regardless of their developmental stage or genotype (Figs. S2H-J). 8 h after exposure, postnatal Nek1-KO and Rad54-KO mice exhibited an increased number of unresolved foci compared to WT mice (Fig. 1D). At this timepoint, most of the mid-S-irradiated cells in Nek1-KO and Rad54-KO mice were in G2 phase, whereas about half of this cell population in WT mice had already entered G1 phase (Fig. 1G). Surprisingly, at neonatal and embryonic stages, Nek1-KO mice had a similar amount of unresolved Rad51 foci as WT mice at 8 h post IR whereas Rad54-KO mice showed considerably more foci (Figs. 1E and 1F). Moreover, the cell cycle progression analysis in Nek1-KO neonatal mice and embryos showed that almost the entire mid-S-irradiated cell population progressed to G1 phase within 8 h after IR, similar to what was observed in WT mice. In Rad54-KO mice, in contrast, the majority of mid-S-irradiated cells remained in G2 phase (Figs. 1H and 1I). The analysis of the small intestines of postnatal and neonatal mice provided similar results (Fig. S3). In conclusion, the analysis of postnatal mice confirms our previous finding that NEK1 regulates the removal of RAD51 foci during HR. Remarkably, however, loss of Nek1 did not affect Rad51 foci removal in tissues from neonatal and embryonic mice, suggesting that Nek1 is dispensable for HR during earlier development.

**Fig. 1:**
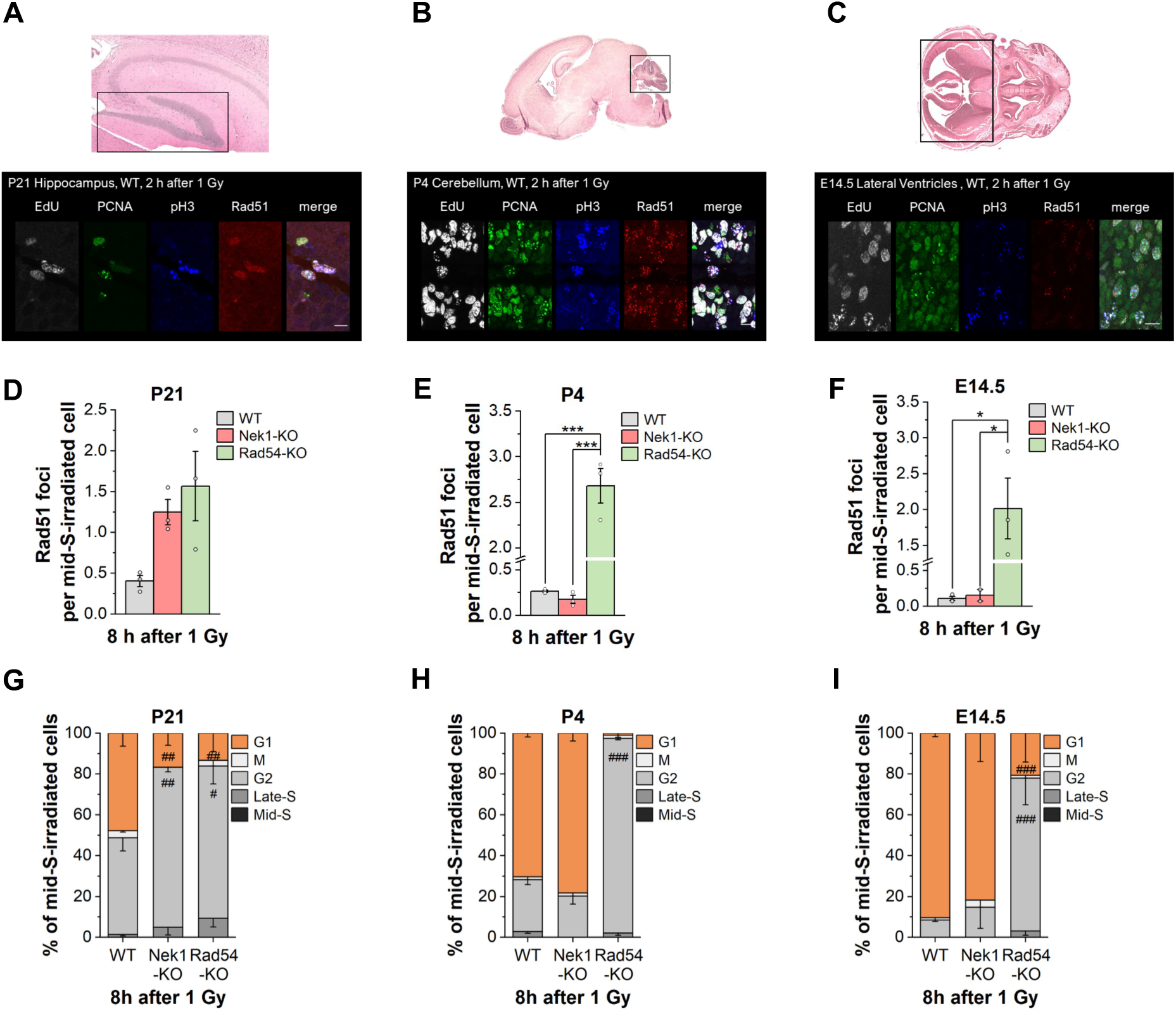
Nek1 is required for HR at late, but not early stages of brain development in vivo. A, B, C: Representative histological (top) and immunofluorescence (bottom) images of brain sections from postnatal (A, P21), neonatal (B, P4) and embryonic (C, E14.5) WT mice. Mice were treated with EdU by intraperitoneal (i.p.) injection, exposed to 1 Gy of X-rays 1.5 h later and sacrificed 2 or 8 h after IR. Brains were isolated, fixed in formaldehyde for 24 h and embedded in paraffin. Sections with a thickness of 4 µm were prepared and stained for cellular structures using hematoxylin and eosin (top) or for cell cycle markers EdU, PCNA, phospho-Histone 3 (pH3) and the HR marker Rad51 using specific kits or antibodies, respectively (bottom). Black boxes indicate areas with proliferative cell populations corresponding to the hippocampus (A), cerebellum (B) and lateral ventricles (C). Scale bars represent 10 µm. D, E, F: Rad51 foci assay in brain tissues of postnatal (D), neonatal (E) and embryonic (F) mice. Mice were treated as described for panel A, B, C. Rad51 foci were quantified 8 h after IR and in all cells that were located in the areas described in A, B, C, and, based on their EdU staining pattern, were irradiated in mid-S phase. G, H, I: Cell cycle analysis in brain tissue of postnatal (G), neonatal (H) and embryonic (I) mice. Mice were treated as described for panel A, B, C. Cycling of irradiated mid-S phase cells was evaluated based on their cellular staining patterns for PCNA (S phase marker) and pH3 (G2 phase marker). Data are shown as mean values of at least 2 animals ± SEM (n≥2). White circles indicate results from individual mice. *,#p < 0.05; **,##p < 0.01; ***,###p < 0.001 (ANOVA).

### Nek1 regulates Rad54 during HR in adult but not in embryonic fibroblasts

Next, we established murine embryonic and postnatal fibroblast lines (MEFs or MPFs, respectively) from WT, Nek1-KO, and Rad54-KO mice and assessed the levels of γH2ax and Rad51 foci after IR. Cells were labeled with EdU, irradiated with 2 Gy, fixed after 2 or 6 h, and stained for cell cycle markers and γH2ax or Rad51. Foci were quantified in EdU-negative G2 phase cells (Figs. S4A-D).^32^ All MPFs exhibited similar numbers of γH2ax foci at 2 h after IR but Nek1-KO and Rad54-KO MPFs showed significantly elevated foci levels compared with WT MPFs at 6 h post IR (Fig. 2A). The same result was obtained for Rad51 with foci numbers being very similar to γH2ax foci numbers at 6 h (Fig. 2C). The corresponding experiments using MEFs provided the same result for Rad54-KO MEFs with elevated γH2ax and Rad51 foci numbers compared with WT MEFs at 6 h post IR. However, Nek1-KO MEFs showed similar γH2ax and Rad51 foci numbers as WT MEFs at 2 and 6 h after IR (Figs. 2B and 2D). We also measured chromatid breaks in G2-irradiated fibroblasts and observed that both Rad54-KO and Nek1-KO MPFs as well as Rad54-KO but not Nek1-KO MEFs exhibited elevated break levels compared with the corresponding WT fibroblasts, fully confirming our analysis of γH2ax and Rad51 foci levels (Figs. S4E-H). These results phenocopy the *in vivo* situation and demonstrate the applicability of the established fibroblast lines for further studies.

**Fig. 2:**
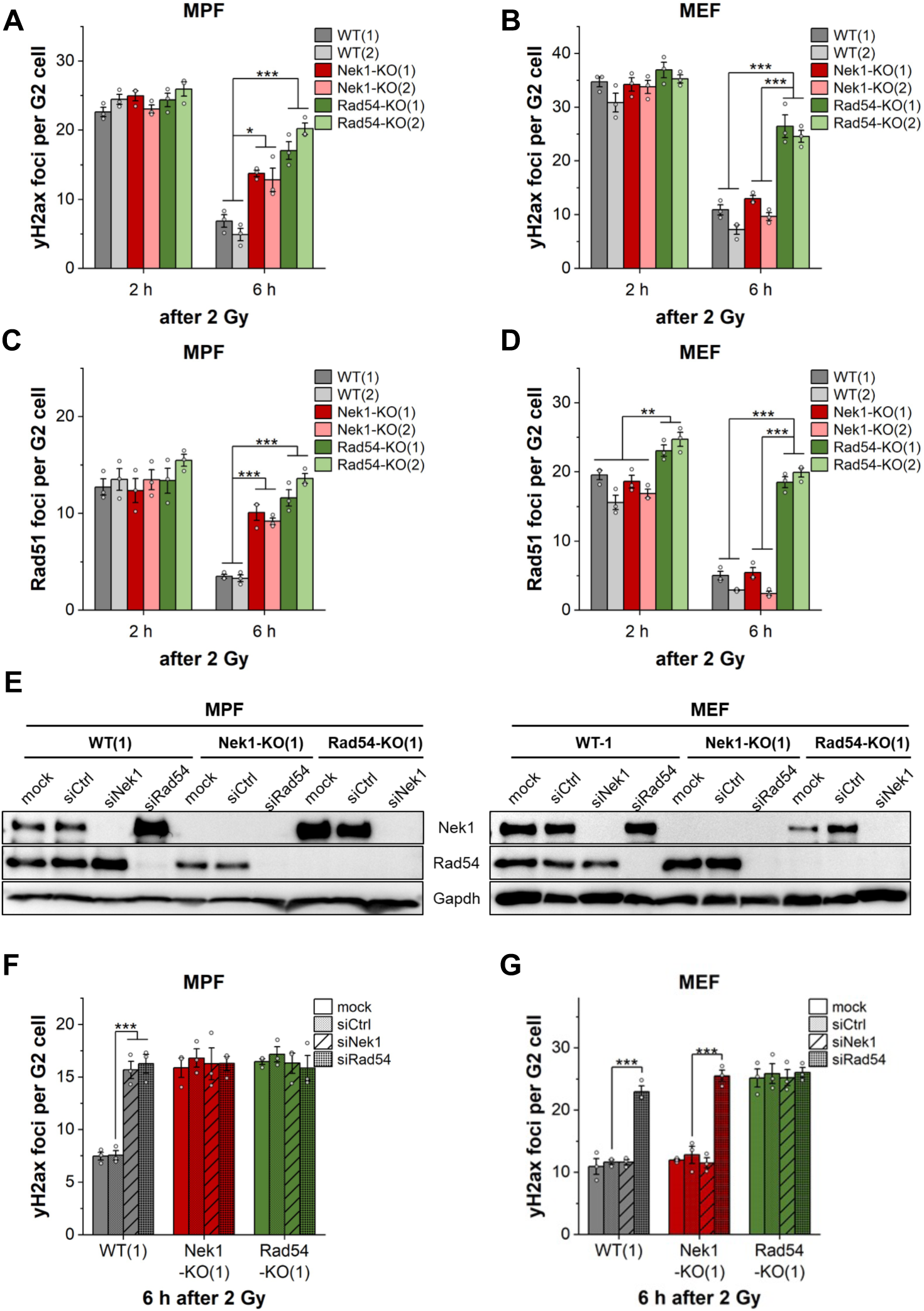
*Nek1 regulates HR in postnatal fibroblasts but is dispensable for HR in embryonic fibroblasts*. A, B: γH2ax foci assay in MPFs (A) and MEFs (B) isolated from two individual postnatal (MPFs) or embryonic (MEFs) WT, Nek1-KO or Rad54-KO mice (1, 2). Fibroblasts were treated with 10 μM EdU for 1 h and irradiated with 2 Gy, fixed 2 h or 6 h thereafter and stained for Rad51, phospho-Histone 3 (pH3) and EdU. C, D: Rad51 foci assay in MPFs (C) and MEFs (D). Fibroblasts were treated as described in A, B but stained for Rad51 instead of γH2ax. E: Representative Western Blots showing the protein levels of Nek1 and Rad54 in WT, Rad54-KO and Nek1-KO MPFs (left) and MEFs (right) following transfection with 100 nM unspecific control, Nek1- or Rad54-specific siRNA for 72 h at standard conditions. Gapdh served as loading control. F, G: γH2ax foci assay in MPFs (F) and MEFs (G). 72 h after transfection with 100 nM unspecific control, Nek1- or Rad54-specific siRNA, fibroblasts were treated with 10 μM EdU for 1 h and then irradiated with 2 Gy, fixed 6 h thereafter and stained as described above. Data are shown as mean values ± SEM (n=3). White circles indicate results from individual experiments derived from 50 EdU-negative, pH3-positive G2 phase cells. Spontaneous foci were subtracted from the irradiated samples. \**p* < 0.05; \*\**p* < 0.01; \*\*\**p* < 0.001 (ANOVA). MPF, murine postnatal fibroblasts; MEF, murine embryonic fibroblasts.

We then depleted Nek1 and Rad54 in MPFs and MEFs by siRNA (Fig. 2E). WT MPFs depleted for either Rad54 (siRad54) or Nek1 (siNek1) exhibited significantly elevated γH2ax foci numbers compared with control siRNA (siCtrl)-treated MPFs (Fig. 2F). Moreover, siRad54 in Nek1-KO MPFs or siNek1 in Rad54-KO MPFs did not further increase γH2ax foci numbers compared with siCtrl-treated Nek1-KO or Rad54-KO MPFs (Fig. 2F), confirming the epistatic relationship of Nek1 and Rad54 previously described. In contrast to MPFs, siNek1 had no effect in WT and Rad54-KO MEF lines while siRad54 caused elevated γH2ax foci numbers in WT and Nek1-KO MEFs (Fig. 2G). siNek1 and siRad54 had no effect in Nek1-KO and Rad54-KO fibroblasts, respectively, confirming the specificity of the siRNAs (Figs. 2F and 2G). Collectively, these data show that the requirement for Nek1 in HR changes during development and raise the possibilities that either Rad54 phosphorylation is dispensable in embryonic cell types or that Rad54 is phosphorylated by other kinases.

### Rad54 phosphorylation is required for HR in both adult and embryonic fibroblasts

Our previous work using HeLa cells suggested that NEK1 phosphorylates RAD54 at S572. To investigate the importance of this phosphorylation in our mouse model, we generated mice with a conditional knock-in (KI) system in exon 12 of the Rad54 gene where S572 is located (Fig. 3A). The KI system consists of two segments, floxed WT Rad54 exons 12-18 connected to the mCherry gene and unfloxed mutant Rad54 exons 12-18 coupled to the EGFP gene. The mutation leads to the substitution of the Ser572 residue to either alanine (Rad54S/A) or glutamate (Rad54S/E). When pairing these mice with Cre mice, the WT exons are removed from the genome through Cre-mediated recombination of the LoxP sites and the mutant Rad54 exons are expressed in the descendant mice. As a result, a GFP-tagged Rad54 protein is produced that, according to our previous study in HeLa cells, cannot be phosphorylated or activated in the Rad54S/A-KI strain or mimics the phosphorylated form and is permanently active in the Rad54S/E-KI strain. Since endogenously produced recombinases can contribute to the formation of DSBs, these mice were backcrossed with WT mice to remove the Cre gene.

**Fig. 3:**
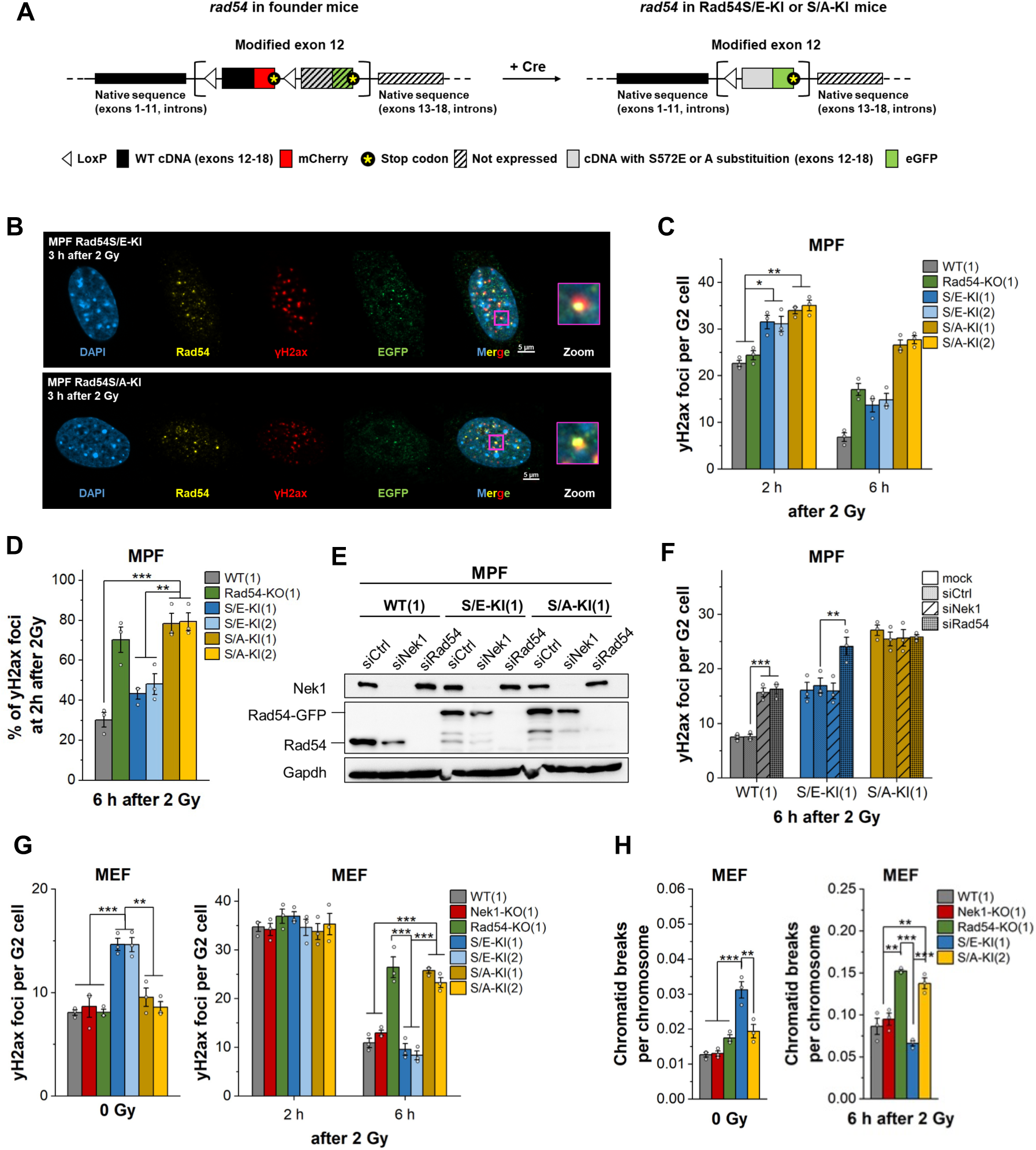
*Functional HR depends on Rad54 phosphorylation in both adult and embryonic fibroblasts*. A: Schematic presentation of the genetic modifications in Rad54-KI mice. In founder mice, Exon 12 of the Rad54 gene was replaced with a conditional knock-in (KI) system which includes two expression cassettes (left). The first cassette consists of floxed WT exons 12-18 and the mCherry gene followed by a stop codon. The second cassette consists of unfloxed Rad54 exons 12 to 18 coupled to GFP with mutations that, at the protein level, substitute the serine at position 572 of Rad54 to either glutamate (S572E) or alanin (S572A). Because of the stop codon behind the mCherry gene, founder mice express an mCherry-tagged WT Rad54 variant. Recombination of the LoxP site flanking the WT Rad54-mCherry cassette was induced by mating these founder mice with mice that express Cre recombinase. As a result, the descendant Rad54S/E-KI or S/A-KI mice express the GFP-tagged Rad54 either containing the S572E or S572A substitution (right). Rad54S/E-KI or S/A-KI mice were finally backcrossed with WT mice to remove the Cre gene from the genome and, thus, to avoid potential Cre-related DSB induction. B: Representative images of colocalization studies in Rad54S/E-KI and S/A-KI MPFs. Fibroblasts were irradiated with 2 Gy, fixed 3 h thereafter and stained for DNA (DAPI, blue), Rad54 (yellow), γH2ax (red), and EGFP (green). Examples of Rad54-GFP-γH2ax colocalizations are depicted in a magnified version (Zoom). Images were taken at a 1000x magnification using an Axioimager M1 microscope equipped with an ApoTome.2. C: γH2ax foci assay in Rad54-KI MPFs isolated from two individual postnatal mice (1, 2). Fibroblasts were treated with 10 μM EdU for 1 h, fixed 2 h or 6 h after IR with 2 Gy and stained for γH2ax, phospho-Histone 3 (pH3) and EdU. Data for WT and Rad54-KO MPFs are taken from Fig. 2A. D: Relative amount of residual γH2ax foci in WT, Rad54-KO and Rad54-KI MPFs. γH2ax foci remaining 6 h post irradiation were normalized to the respective foci numbers 2 h after irradiation. E: Representative Western Blot showing the protein levels of Nek1 and Rad54 in Rad54-KI MPFs following transfection with 100 nM unspecific control, Nek1- or Rad54-specific siRNA for 72 h at standard conditions. Gapdh served as loading control. F: γH2ax foci assay in Nek1-and Rad54-depleted Rad54-KI MPFs. 72 h after transfection with 100 nM unspecific control, Nek1- or Rad54-specific siRNA, fibroblasts were treated with 10 µM EdU for 1 h and then irradiated with 2 Gy, fixed 6 h thereafter and stained for γH2ax, phospho-Histone 3 (pH3) and EdU. G: γH2ax foci assay in Rad54-KI MEFs isolated from two individual postnatal embryonic mice (1, 2). Fibroblasts were treated with 10 μM EdU for 1 h, fixed 2 h or 6 h after IR with 2 Gy, or left unirradiated, and stained for γH2ax, phospho-Histone 3 (pH3) and EdU. Data for WT(1), Nek1-KO(1) and Rad54-KO(1) MEFs are taken from Fig. 2B. H: Chromatid breaks in WT, Nek1-KO, Rad54-KO and Rad54-KI MEFs. Fibroblasts were fixed 6 h after IR with 2 Gy or left unirradiated and chromosome spreads were obtained from G2-phase cells. Data are shown as mean values ± SEM (n=3). White circles indicate results from individual experiments derived from 50 EdU-negative, pH3-positive G2 phase cells or at least 40 chromosome spreads. Spontaneous foci and chromatid breaks were subtracted from the irradiated samples. *p < 0.05; **p < 0.01; ***p < 0.001 (ANOVA). MPF, murine postnatal fibroblasts; MEF, murine embryonic fibroblasts.

For the following studies, postnatal fibroblast lines were established from both Rad54-KI strains, spontaneously immortalized, and genotyped by PCR to verify homozygosity of the KI alleles and absence of the WT Rad54 and Cre genes. The functional expression of both Rad54 mutants was controlled with immunofluorescence assays. For this, Rad54-KI cells were irradiated with 2 Gy, fixed 3 h later, and stained for Rad54, GFP, and either mCherry or the DSB marker γH2ax. Rad54S/A-KI and Rad54S/E-KI cells exhibited Rad54 foci which colocalized with GFP but not mCherry, confirming both the expression of the Rad54 mutants and the removal of the WT Rad54 gene (Fig. S5A). Moreover, both proteins colocalized with γH2ax foci indicating that the introduced mutations did not interfere with the recruitment of Rad54 to sites of DNA damage (Fig. 3B). Finally, we performed γH2ax foci repair measurements as described above for KO cells to determine how the modified Rad54 proteins affect HR. Unirradiated Rad54S/E-KI cells accumulated twice as many spontaneous γH2ax foci as WT, Rad54-KO or Rad54S/A-KI postnatal cells (Fig. S5D), suggesting that constitutively active Rad54 enhances endogenous replication stress.^20^ At 2 h after IR, WT, Rad54-KO and Rad54-KI cells exhibited slightly different γH2ax foci levels (Fig. 3C), possibly because of differences in DNA content of the spontaneously transformed cell lines. To correct for this effect, γH2ax foci numbers at 6 h were normalized to foci numbers at 2 h. Rad54S/E-KI cells showed a level of γH2ax foci similar to WT cells, indicating their proficiency to repair DSBs. Rad54S/A-KI cells, in contrast, exhibited elevated γH2ax foci numbers similar to Rad54-KO cells (Fig. 3D). These data demonstrate the importance of the S572 residue for the regulation and activation of Rad54 in postnatal fibroblasts.

To verify the S572 residue as the site for Nek1-controlled phosphorylation, γH2ax foci were analyzed in Rad54-KI MPFs transiently depleted for Nek1 and Rad54 (Fig. 3E). The level of γH2ax foci at 6 h post IR in Rad54S/E-KI MPFs was not enhanced by siNek1 while siRad54 caused a significant increase, indicating that the permanently active form of Rad54 rescues the DSB repair defect observed in siNek1-treated WT MPFs (Fig. 3F). Depletion of Nek1 or Rad54 in Rad54S/A-KI cells did not further enhance the already elevated γH2ax foci level, showing that Nek1 deficiency exerts no additional repair defect beyond that caused by the absence of S572 phosphorylation (Fig. 3F). Consistent with the foci analysis, we also observed increased levels of chromatid breaks in siRad54-treated but not in siNek1-treated Rad54S/E-KI MPFs at 6 h post IR. In contrast, both siRad54 and siNek1 increased break levels in WT MPFs and neither siRad54 nor siNek1 affected break levels in Rad54S/A-KI MPFs at 6 h post IR (Fig. S5E). Additionally, Rad54S/E-KI MPFs exhibited substantially higher levels of spontaneous chromatid breaks which were reduced after depletion of Rad54 but not Nek1 (Fig. S5E). These data demonstrate that murine Nek1 mediates, either indirectly or directly, the phosphorylation of Rad54 at S572 and promotes HR in postnatal fibroblasts, fully confirming our previous work with HeLa cells.^20^

In contrast to postnatal fibroblasts, depletion of Nek1 did not diminish the efficiency of HR in MEFs (see Fig. 2G). To explore if Rad54 phosphorylation is dispensable for HR in embryonic stages, we isolated MEFs from both Rad54-KI mouse strains and verified their functionality (Figs. S5B and S5C). MEFs expressing constitutively active Rad54S/E accumulated high levels of spontaneous γH2ax foci but were able to efficiently repair IR-induced γH2ax foci (Fig. 3G). MEFs expressing phospho-defective Rad54S/A, in contrast, exhibited normal spontaneous but elevated IR-induced γH2ax foci levels similar to Rad54-KO cells (Fig. 3G). Additionally, Rad54S/E but not Rad54S/A MEFs showed high levels of spontaneous chromatid breaks and Rad54S/A but not Rad54S/E MEFs failed to repair IR-induced chromatid breaks (Fig. 3H), recapitulating the findings of our foci analysis. The elevated spontaneous break levels in constitutively active Rad54S/E postnatal and embryonic fibroblasts shows that Rad54 requires regulation by phosphorylation in both developmental stages. The failure of Rad54S/A MEFs to repair IR-induced breaks demonstrates that Rad54 phosphorylation is important for HR in embryonic cells, as it is in postnatal cells. However, since Nek1 is completely dispensable in MEFs (Figs. 3G and 3H), kinases other than Nek1 are required to phosphorylate Rad54 during early developmental stages.

### Nek3 and Nek5 regulate Rad54 phosphorylation during HR in embryonic fibroblasts

We next explored if another member of the Nek kinase family might phosphorylate Rad54 in embryonic cells. First, we analyzed the expression levels of all Nek genes in WT MEFs relative to WT MPFs using a SYBR green-based qPCR assay. *Nek1*, *2*, *6*, *7*, *8*, *9*, *10*, and *11* were similarly expressed in both MEFs and MPFs (Fig. 4A). *Nek4* and *Nek10* were significantly downregulated in MEFs compared to MPFs whereas *Nek3* and *Nek5* were expressed 3-fold and 5.5-fold higher in MEFs (Fig. 4A), possibly suggesting a role for these kinases during embryonic stages.

**Fig. 4:**
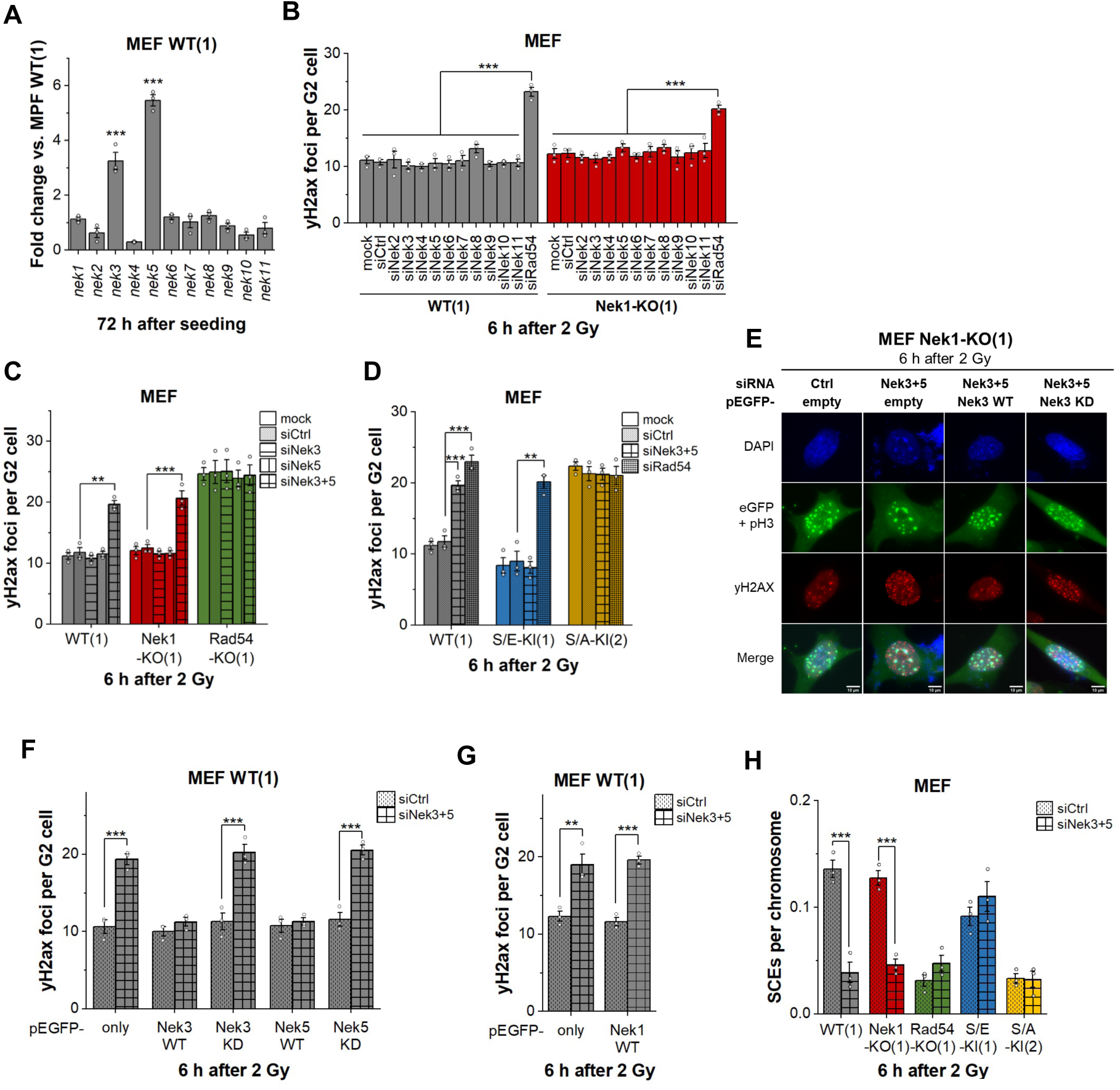
*Nek3 and Nek5, but not Nek1, regulate HR in embryonic fibroblasts*. A: Differential gene expression analysis in untreated WT MEFs. RNA was isolated from untreated fibroblasts 72 h post seeding, converted into cDNA, and finally analyzed using qPCR and specific primers. Fold changes in expression were calculated with the 2^-ΔΔCt^ method using Gapdh as a reference gene and normalizing the expression levels of each Nek gene in WT MEFs to their corresponding level in WT MPFs. B: γH2ax foci assay in WT and Nek1-KO MEFs depleted for other Nek family members. 72 h after transfection with unspecific control or Nek-specific siRNAs, fibroblasts were treated with 10 µM EdU for 1 h and then irradiated with 2 Gy, fixed 6 h thereafter, and stained for γH2ax, phospho-Histone 3 (pH3), and EdU. C: γH2ax foci assay in WT, Nek1-KO, Rad54-KO MEFs following transfection with 100 nM unspecific control or Nek3- and Nek5-specific siRNA for 72 h at standard conditions and treatment as described for panel B. D: γH2ax foci assay in Rad54-KI MEFs following transfection with 100 nM unspecific control, Rad54- or Nek3- and Nek5-specific siRNA for 72 h at standard conditions and treatment as described in panel B. Data for WT(1) are taken from panel C. E: Representative fluorescence images of the rescue assay in WT and Nek1-KO MEFs. Fibroblasts were treated with 100 nM unspecific control or Nek3- and Nek5-specific siRNAs for 24 h at standard conditions followed by transfection with 10 µg of plasmids coding for only GFP, the wildtype form of Nek3 (WT) or a kinase-dead variant of Nek3 (KD). 48 h after plasmid transfection, fibroblasts were treated with 10 µM EdU for 1 h and then irradiated with 2 Gy, fixed 6 h thereafter and stained for γH2ax, phospho-Histone 3 (pH3), GFP and EdU. F, G: Quantification of rescue assays in WT MEF as shown in panel E. γH2ax foci were quantified in 50 EdU-negative, pH3/GFP-positive G2-phase cells. H: SCE assay in WT, Nek1-KO, Rad54-KO and Rad54-KI MEFs following transfection with 100 nM unspecific control or Nek3- and Nek5-specific siRNA for 72 h at standard conditions. Fibroblasts were fixed 6 h after IR with 2 Gy and chromosome spreads were obtained from mitotic cells. Data are shown as mean values ± SEM (n=3). White circles indicate results from individual experiments which in case of γH2ax foci assays shown in panel B, C and D were derived from 50 EdU-negative, pH3-positive G2 phase cells or in case of SCE measurements shown in panel H were derived from at least 40 chromosome spreads. Spontaneous foci and SCEs were subtracted from the irradiated samples. \**p* < 0.05, \*\**p* < 0.01; \*\*\**p* < 0.001; versus WT MPFs or siCtrl (ANOVA). MPF, murine postnatal fibroblasts; MEF, murine embryonic fibroblasts.

We then performed a functional siRNA screen to evaluate the effect of Nek depletion on HR. WT MEFs were treated with Nek-specific siRNAs and analyzed using the γH2ax foci repair assay; siRNA efficiencies were validated using the qPCR assay (Fig. S6A). In contrast to the loss of Rad54, the depletion of any of the Nek members did not elevate γH2ax foci numbers compared with siCtrl-treated WT MEFs (Fig. 4B). We also analyzed Nek1-KO MEFs and obtained the same result, excluding the possibility that Nek1 and any of the other family members function redundantly during HR in MEFs (Fig. 4B). Finally, we tested if depletion of a combination of Nek kinases causes elevated unrepaired γH2ax foci in MEFs. Since Nek3 and Nek5 are phylogenetically closer related to Nek1 than the other Nek family members and are also significantly higher expressed in MEFs compared to MPFs, we combined the depletion of Nek3 and Nek5 (Fig. S6B) and assessed γH2ax foci levels in WT and Nek1-KO MEFs. Strikingly, the combined depletion of Nek3 and Nek5 caused significantly elevated foci levels compared with siCtrl-treated WT and Nek1-KO MEFs (Fig. 4C). In contrast, the combined depletion of Nek3 and Nek5 did not further elevate γH2ax foci levels in Rad54-KO MEFs (Fig. 4C), demonstrating that Nek3/5 function in MEFs via the Rad54 pathway. Nek1 does not exert a role during HR in MEFs as WT and Nek1-KO MEFs are similarly affected by the depletion of Nek3 and Nek5. Finally, Nek3 and Nek5 do not function during HR in MPFs since the single depletion of Nek1 in WT MPFs causes a repair defect epistatic to Rad54 whereas the combined depletion of Nek3 and Nek5 in WT MPFs has no effect (Figs. S6C and S6D).

To determine whether Nek3 and Nek5 regulate HR in embryonic cells similarly to Nek1 in postnatal cells by mediating the phosphorylation of Rad54 at Ser572, Nek3/5-depleted Rad54-KI MEFs were analyzed with the γH2ax foci repair assay. Strikingly, the combined Nek3 and Nek5 depletion in Rad54S/E-KI MEFs did not increase γH2ax foci numbers compared with siCtrl-treated Rad54S/E-KI MEFs (Figs. 4D and S6B), indicating that Nek3 and Nek5 are dispensable in cells with constitutively active Rad54. Moreover, siNek3/5 did not further increase γH2ax foci numbers in Rad54S/A-KI MEFs (Fig. 4D), confirming that they function via the Rad54 pathway. Collectively, these data demonstrate that HR in embryonic cells depends on Nek3 and Nek5 for the activation of Rad54.

Finally, we wished to confirm the redundant roles of Nek3 and Nek5 during HR in MEFs using rescue experiments. We depleted Nek3/5 in WT and Nek1-KO MEFs, transfected the cells with plasmids expressing siRNA-resistant, GFP-tagged Nek3 or Nek5 and analyzed γH2ax foci in GFP-positive G2 cells (Fig. 4E). Depletion of Nek3/5 in WT and Nek1-KO MEFs expressing a GFP-only plasmid caused elevated γH2ax foci numbers compared with siCtrl-treated WT and Nek1-KO MEFs while the expression of a siRNA-resistant, GFP-tagged Nek3 or Nek5 plasmid rescued this defect (Figs. 4F and S6F). As a control, we expressed a GFP-tagged Nek1 plasmid in Nek3/5-depleted MEFs and did not observe a rescue of the repair defect (Fig. 4G). Similarly, expression of a GFP-tagged Nek1 plasmid but not a Nek3 or Nek5 plasmid did rescue the repair defect of Nek1-KO MPFs (Fig. S6G), excluding the possibility that these plasmid expression experiments overrule the specific requirements of the Nek family members during development. Collectively, these experiments confirm that either of the two kinases alone, Nek3 or Nek5, can mediate the phosphorylation of Rad54 in MEFs. Finally, we explored if the kinase functions of Nek3 and Nek5 are required for mediating Rad54 phosphorylation and generated kinase-dead mutants. Similar to the already published kinase-dead form of Nek1, we substituted the highly conserved Lysin33 residue in Nek3 and Nek5 with Arginine (K33R). Of note, neither the K33R mutant of Nek3 nor of Nek5 was able to rescue the elevated γH2ax foci level of siNek3/5-depleted WT or Nek1-KO MEFs (Fig. 4F). Thus, both Nek3 and Nek5 function as regulatory kinases during HR in MEFs.

Since Nek3 and Nek5 have not been previously implicated in HR repair of DSBs, we wished to confirm their described role by using an assay directly assessing HR efficiency. For this, we irradiated cells in G2 phase and analyzed the formation of sister chromatid exchanges (SCEs) (Fig. S6H). SCEs after IR arise exclusively from the repair of DSBs by HR and Rad54 has been shown to promote their formation.^20,33,34^ We observed approximately 0.15 IR-induced SCEs per chromosome at 6 h post 2 Gy in siCtrl-treated WT and Nek1-KO MEFs (about 8 SCEs per cell) and a strong reduction after the dual depletion of Nek3 and 5 (Fig. 4H and S6I) (note that about 8 SCEs arise from the repair of about 16 Rad51 foci, see Fig. 2D). Importantly, Rad54-KO and Rad54S/A-KI MEFs showed substantially fewer SCEs which were not further diminished by siNek3/5 while Rad54S/E-KI MEFs showed SCEs levels similar to WT and Nek1-KO MEFs and siNek3/5 did not reduce this level (Figs. 4H and S6I). This analysis establishes Nek3 and Nek5 as HR repair factors in embryonic cells and confirms that they function by mediating the phosphorylation of Rad54 at S572.

## Discussion

### Nek1 mediates Rad54 phosphorylation and HR during later developmental stages

HR represents an important pathway for repairing DSBs but HR factors, including RAD51, also serve to protect and restart stalled replication forks. RAD54 functions during DSB repair where it removes RAD51 from heteroduplex DNA which is formed during D-loop formation.^35^ This allows subsequent DNA repair synthesis and completion of the HR process. We have previously suggested that RAD54’s activity is regulated by NEK1 in a cell cycle-specific manner to promote RAD51 removal from dsDNA and HR in G2 phase without interfering with RAD51’s fork stabilization role during S phase which requires binding to dsDNA.^18–20^ Here, we provide evidence using a mouse model that Nek1 exerts its HR function during later developmental stages. Further, we show that Nek1 is dispensable for HR during embryogenesis and that Nek3 and Nek5 activate Rad54 during this developmental stage.

The cellular role of NEK1 has raised particular interest since it was discovered that NEK1 is frequently mutated in patients with ALS.^36,37^ Of note, another frequently mutated ALS factor, C21ORF2, controls the cellular function of NEK1 and its loss has been shown to compromise HR similar to the loss of NEK1.^38^ However, the role of NEK1, and C21ORF2, remained unclear since the study did not observe a defect in removing IR-induced RAD51 foci in cells deficient for NEK1 and C21ORF2.^38^ Here, we addressed this in the following ways. First, we analyzed Rad51 foci removal *in vivo* in various tissues of Nek1-KO mice in comparison to WT and Rad54-KO mice and *in vitro* in fibroblasts derived from these mice. We used a cell cycle-specific approach and demonstrate the role of Nek1 in Rad51 foci removal for adult, but not for embryonic, mice as well as for fibroblasts derived from adult mice but not for fibroblasts derived from embryonic mice. Second, we generated Rad54-KI mice where the endogenous S572 locus is mutated to either unphosphorylatable alanine (S/A) or phospho-mimic glutamate (S/E). Rad54S/A-KI fibroblasts derived from adult mice show a defect in the repair of IR-induced γH2ax foci similar to the defect observed in fibroblasts derived from adult Nek1-KO or Rad54-KO mice. Importantly, phospho-mimic Rad54S/E-KI fibroblasts are proficient in repairing IR-induced γH2ax foci even in the absence of Nek1. Collectively, our approaches demonstrate that Nek1 exerts its pro HR function during later developmental stages by mediating the phosphorylation of Rad54 at S572.

### Deregulated Rad54 activity leads to replication stress

Our evaluation of spontaneous γH2ax foci levels in embryonic as well as adult Rad54-KI cells and our previous work including the analysis of replication fork stability using the DNA fiber assay in HeLa cells collectively suggest that constitutively active Rad54S/E interferes with Rad51’s function in stabilizing stalled replication forks.^20^ Moreover, our previous analysis using a phospho-specific antibody showed that RAD54-S572 phosphorylation takes place in G2 phase even if DNA damage is induced during S phase. Thus, to minimize replication stress and promote efficient HR, RAD54 activation needs to be regulated in a cell cycle-specific manner such that its activity is restricted to G2 phase. This suggested cell cycle regulation of RAD54 activity is consistent with studies where the repair of daughter strand gaps involves RAD51 loading at the replication fork but completion of the process uncoupled from replication in the following G2 phase.^14,39^ NEK family members might be particularly suitable to mediate the cell cycle-specific phosphorylation of RAD54 since their other roles in ciliogenesis, microtubule and centromere organization as well as checkpoint regulation also demand a cell cycle-specific activity.

### Nek3 and Nek5 mediate Rad54 phosphorylation and HR in mouse embryonic fibroblasts

NEK3 and NEK5 have no established roles in the DNA damage response and very little information is available about their functions. A yeast two-hybrid screen identified 27 potential NEK3 interactors, including PCNA which functions during DNA replication and repair, but the functional relevance of this interaction was not studied.^40^ NEK5 was reported to be involved in G2 checkpoint regulation and repair of etoposide-induced DNA damage assessed by the alkaline comet assay.^41^ Interestingly, an extensive phylogenetic analysis has described NEK1, NEK3 and NEK5 as the only members of one of the five identified NEK subfamilies.^42^ Here, we show that both kinases, Nek3 and Nek5, function redundantly to mediate Rad54 phosphorylation and promote HR in murine embryonic fibroblasts. Similar to the other Nek family members, they have cell cycle regulatory functions which make them suitable for coordinating the process of HR with progression from S to G2 phase. Since the cell cycle-specific Nek activities are likely to change during development, a coordinated handover from one to another family member might be needed to maintain the strict cell cycle requirement for Rad54 phosphorylation. Moreover, the duration of the different cell cycle phases likely changes during development, with embryonic cells typically exhibiting a longer S phase, which might demand a switch in the kinase requirement for G2-specific Rad54 phosphorylation.^43^ Since our analysis was limited to fibroblasts from two developmental stages, embryonic stage E14.5 and adult stage P21, it is possible that additional Nek family members are involved in regulating HR during other developmental stages. Our observation that Nek3 and Nek5 can both mediate Rad54 phosphorylation might suggest that embryonic stage 14.5 represents a handover period from one to the other of these two kinases during which they are both active. Alternatively, the combined activity of Nek3 and Nek5 might provide a more ideal condition to coordinate Rad54 phosphorylation with cell cycle progression. Collectively, we not only identified two new kinases involved in DSB repair by HR but also report a hitherto unknown developmental change in the regulation of HR.

### A possible connection to ALS

Mutated NEK1 is linked to the neurogenerative disease amyotrophic lateral sclerosis (ALS) which affects the viability of motor neurons and is usually diagnosed at an age between 50 and 60.^36,44^ Although the exact cause of the disease remains elusive, early research has already suggested a mechanism related to dysfunctions in the DDR since ALS-derived motor neurons accumulate a large amount of DNA damage and chromosomal aberrations.^45,46^ Genetic studies of patients with a familial history of ALS revealed mutations in about 30 genes mostly related to autophagy, the antioxidative system and the DDR.^47,48^ Two independent mutations of NEK1 resulting in a loss-of-function and a missense mutation have been identified as risk factors and are associated with 3% of all spontaneous and familial ALS cases.^49,50^

ALS and other neurodegenerative and neurodevelopmental diseases have been suggested to represent pathological outcomes of replication stress and subsequent death of specific neuronal cell types.^51^ Many neuron-specific genes are very large and have an increased risk of replication-transcription conflicts, a known cause of replication stress.^52,53^ HR serves to limit damage from replication stress by repairing DSBs that arise at broken replication forks.^54,55^ Damage to neuronal genes is further minimized by restricting their transcription to terminally differentiated cells.^56^ However, changes in the control of transcription, replication and cell cycle regulation will cause replication stress in neuronal genes and the miscoordination with differentiation will likely enhance the damage load. With this conceptual framework in mind, it might be expected that cellular regulators with roles in several of these pathways are particularly prone to cause neurodegenerative diseases if mutated. Of note, differentiation of neuronal cells is not limited to early developmental stages but can occur throughout adulthood.

NEK family members in general are key regulators of cell cycle-specific processes associated with development and differentiation.^24^ Nek1 and Nek3/5 are, as we show, additionally involved in HR by regulating the phosphorylation of Rad54. Notably, mis-regulated Rad54 can cause enhanced levels of replication stress exemplified by increased spontaneous γH2ax foci levels. Thus, mutations in Nek family members have not only the potential to affect the differentiation process but could also enhance the level of replication stress and/or alter the efficiency of HR which is needed to counteract DNA damage from replication stress. It is, therefore, perhaps not surprising that NEK1 is one of the most frequently mutated genes in familial as well as sporadic forms of ALS. Moreover, it has not escaped our attention that also NEK5 has been found to be mutated in several ALS patients, although at a lower frequency compared with NEK1.^47^ Finally, an association between neurodegenerative diseases and the mis-regulation of HR has also been described for mutations in other factors, including ATRX, CHAMP1, ATM, ATR.^57–60^

## Material and Methods

### Mouse models

WT mice (C57BL/6, Strain#:000664), Nek1-KO mice (C57BL/6 Nek1^kat-2J^, Strain#: 002854), and Cre mice (B6.C-Tg(CMV-cre)1Cgn/J, Strain#: 006054) were purchased from The Jackson Laboratory, Maine, USA. Rad54-KO mice (C57BL/6 mRad54^307neo^) were kindly provided by Prof. Dr. Kanaar from the Erasmus Medical Center in the Netherlands.^61^ Founder mice for the Rad54S/E-KI and Rad54S/A-KI strains were produced by Cyagen, USA, and bred in-house with Cre mice to generate offspring expressing one of the mutant Rad54 variants. Successful Cre-mediated deletion of floxed genes was confirmed by PCR. Offspring were subsequently crossed with WT mice to remove the Cre gene. All mice were bred in-house according to institutional and national guidelines for laboratory animals. Breeding of KI mice was approved by the responsible committee (Regierungspräsidium Darmstadt, Darmstadt, Germany, DA8/1009) and conducted in accordance with the requirements of the German Animal Welfare Act.

### In vivo Rad51 foci and cell cycle progression assay

Experimental mice were bred in-house and confirmed for homozygosity of the KO or KI genes by PCR. WT mice used in the experiments were derived from pairings of heterozygous Nek1-KO or Rad54-KO mice. Female and male mice were included in the study. For DNA repair studies in the embryonic brain, female mice were checked daily after mating for the presence of a vaginal plug. 14.5 days after the plug occurred, pregnant mice were treated with 38 µg of 5-ethynyl-2’-deoxyuridine (EdU, Roth, Germany) dissolved in sterile 0.9% saline solution per gram body weight by intraperitoneal (i.p.) injection. EdU distributes systemically and is integrated into the DNA of proliferating cells during replication in S phase. 1.5 h after EdU injection, mice were similarly treated with 76 µg/g 5-bromo-2’-deoxyuridine (BrdU, Sigma-Aldrich), a second thymidine analogue which is incorporated into DNA preferentially over EdU. This treatment plan made it possible to identify proliferating cells in various tissues and discriminate between cells that were in S phase at the time of irradiation (EdU positive) or entered S phase after irradiation (BrdU positive). Immediately after BrdU injection, mice were irradiated with an X-RAD320 X-ray tube (Precision, USA) at 250 kV using a 1 mm aluminum filter and a dose rate of ∼1 Gy/min. The same procedure was performed on neonatal and postnatal mice, which were analyzed at 4 days (P4) or 21 days (P21) after birth, respectively. At the time points indicated in the figures, mice were sacrificed, and tissues were fixed for 24 h in 4% neutral-buffered formalin, except for the brains of P21 mice, which were fixed in 10% formalin. Tissues were then embedded in paraffin and sliced into sections with a thickness of 4 µm. All *in vivo* experiments were approved by the responsible committee (Regierungspräsidium Darmstadt, Darmstadt, Germany, DA8/1003) and conducted in accordance with the requirements of the German Animal Welfare Act.

### Immunohistochemistry and imaging

Tissue sections were dewaxed, rehydrated, and treated with 1x Universal HIER antigen retrieval solution (Abcam, UK) for 35 min at 98°C. Samples were incubated with primary antibodies diluted in 0.2% Triton-X100/PBS at 4°C overnight and with secondary antibodies for 2-4 h at 37°C in the dark after washing with prewarmed 0.1% Tween20/PBS (10 mM Na₂HPO₄, 1.8 mM KH₂PO₄, 137 mM NaCl, 2.7 mM KCl). If primary antibodies from the same species were used, the staining protocol described above was performed twice in a row with an additional wash step between the first secondary and the second primary antibody. Antibodies included: Mouse anti-Nek1 (Santa Cruz Biotechnology, 1:200), Goat anti-Rad54 (Santa Cruz Biotechnology, 1:100), Mouse anti-BrdU (DSHB, 1:500), rabbit anti-Rad51 antibody (Calbiochem, 1:300), Mouse anti-PCNA (Santa Cruz Biotechnology, 1:600), Rabbit anti-phospho-Histone 3 (pH3, Merck, 1:300), Donkey anti-mouse AlexaFluor488 (Invitrogen, 1:600), Donkey anti-mouse DyLight550 (Abcam, 1:600), Donkey anti-rabbit DyLight405 (Dianova, 1:600), Donkey anti-rabbit DyLight550 (ThermoFisher Scientific, 1:600) and Donkey anti-goat AlexaFluor488 (Molecular Probes, 1:600). EdU was stained using the Click-iT EdU Imaging Kit (Baseclick, Germany) as instructed by the manufacturer before staining DNA with 0.1 µg/ml DAPI (Sigma-Aldrich, Germany). Slides were mounted in Vectashield mounting medium (Axxora, Germany) and sealed with nail polish. Images from immunostained tissues were taken on a confocal microscope (TCS SP5 II, Leica, Germany) with LAS AF Lite software (Leica, Germany) at 40x, 63x, or 100x magnification. Images were taken as maximum intensity projections of stacks (z = 5-20). Images were processed and analyzed using Fiji (ImageJ, version 1.54i). Cell cycle progression was measured by determining the pH3 and PCNA signals in at least 40 cells that showed a specific EdU signal pattern corresponding to early S, mid S or late S phase. In the same cells, Rad51 foci were quantified. H&E stainings were performed by incubating sections in Hematoxylin solution (Roth, Germany) for 10 min at room temperature (RT) and in EosinG solution (Merck, Germany) for 10 s, followed by dehydration, mounting in Coverquick 4000 (Labonord, France), and sealing. H&E images were captured using an Axioimager M1 microscope with ApoTome.2 (Zeiss, Germany) and Metafer4 (MetaSystems) software at 10x and 40x magnification and arranged with Fiji.

### Cloning and mutagenesis of Nek1, Nek3, and Nek5 vectors

Murine cDNA of Nek3 or Nek5 was cloned from pCMV-SPORT6 plasmids (Dharmacon, USA) into pEGFP-C1 plasmids (Addgene, USA) using XmaI and XbaI (New England Biolabs, USA). Vectors were amplified in *E. coli* DH10b or DH5a cells (Thermo Fisher Scientific, Germany) before cloning. *E. coli* cells were mixed with 20 ng plasmid DNA and transformed using the heat shock method: 30 min on ice, 45 s at 42°C, and 2 min on ice. Cells were then resuspended in SOC medium (Thermo Fisher Scientific, Germany) and incubated for 1 h at 37°C, followed by plating onto LB plates (2.5% LB medium, 1.5% Agar-Agar) supplemented with appropriate antibiotics. After overnight incubation at 37°C, single colonies were transferred into LB medium (2.5% LB medium) supplemented with appropriate antibiotics and cultured overnight at 37°C. Plasmids were isolated using the Z-R Plasmid Miniprep Classic Kit (Zymo Research, USA) or the Genopure Plasmid Maxi Kit (Roche) as instructed by the manufacturers, and 1 µg of plasmid was digested with 10 units of XmaI and 10 units of XbaI overnight at 37°C. The digested DNA was separated on an agarose gel (1.5% agarose, 0.6 µl Roti-Safe gel stain/ml gel) in 0.5x TBE (45 mM Tris pH 8.4, 45 mM Boric acid, 1 mM EDTA), and fragments of interest were isolated using the QIAquick Gel Extraction Kit (Qiagen, Germany) according to the manufacturer’s instructions. The linearized pEGFP plasmid was incubated with Nek3 or Nek5 cDNA fragments and T4 DNA Ligase in ligase buffer as instructed by the manufacturer (New England Biolabs, USA). Correct ligation was confirmed with a control digest of the pEGFP constructs with XhoI and NotI-HF (New England Biolabs, USA) and sequencing with primers available at Microsynth AG (EGFPC-for, SV40-pArev). Murine cDNA of Nek1 was cloned from pCMV6-Entry plasmids (Origene, USA) into pEGFP-C1 plasmids (Addgene, USA) using SalI-HF and MluI-HF (New England Biolabs, USA). Site-directed mutagenesis (SDM) was applied to confer siRNA resistance and to generate kinase-dead (K33R) variants of Nek1, Nek3 and Nek5. According to a protocol by Edelheit *et al.*, two samples per pEGFP plasmid were set up with 500 ng DNA and 40 pmol of either the forward or reverse primer.^62^ The following primers (forward, reverse, 5’-3’) were used: siRNA resistance Nek3 (aag ata acg aaa acc ctg att ggc taa gcg aac taa aga agc acg tag gat acg, cgt atc cta cgt gct tct tta gtt cgc tta gcc aat cag ggt ttt cgt tat ctt); siRNA resistance Nek5 (agt ttc agg agc aca gat gta agg agg aac acg agg att aca cag aca gag cct ttg, caa agg ctc tgt ctg tgt aat cct cgt gtt cct cct tac atc tgt gct cct gaa act); K33R-substitution Nek1 (ttg aga tgt taa ttt ccc tga tga cat aat gtc tgc cat cctc, gag gat ggc aga cat tat gtc atc agg gaa att aac atc tcaa); K33R-substitution Nek3 (gca atc aga cat ttg cca tga ggg aaa tca gac tgctc, gag cag tct gat ttc cct cat ggc aaa tgt ctg attgc); K33R-substitution Nek5 (cag aaa gca gtc act gtg tca taa gag aaa tca gtt tga caaag, ctt tgt caa act gat ttc tct tat gac aca gtg act gct ttctg). Samples were supplemented with Q5 polymerase (New England Biolabs, USA) and amplified in a thermal cycler under the following conditions: 94°C for 2 min, 30 cycles of 94°C for 40 s, 55°C for 40 s, and 72°C for 1 min/kb, followed by a final step at 72°C for 7.5 min. Combined reverse and forward amplified samples were cooked at 95°C for 5 min, cooled to 37°C and supplemented with 30 units of the restriction enzyme DpnI (New England Biolabs, USA). The resulting plasmids were amplified in *E. coli* DH5α cells and analyzed as described above.

### Fibroblast isolation and culture

Fibroblasts were established from WT and homozygous KO and KI mice. For the generation of murine embryonic fibroblast (MEF) lines, female mice were checked daily after mating for the presence of a vaginal plug. Embryos were harvested from pregnant mice 13.5 days post fertilization, following the protocol by Durkin *et al.*^63^ The isolated embryos were decapitated and digested in accutase solution (Sigma-Aldrich, Germany) at 37°C. Murine postnatal fibroblast (MPF) lines were prepared from the ears of 21-day-old mice, as described by Khan *et al.*^64^ Mice were sacrificed and their ears were excised, disinfected in 70% ethanol (Roth, Germany), and then digested in a Pronase E/Collagenase D enzyme mixture (1.25 µg/µl Pronase E in 1 mM EDTA, 10 mM Tris pH 8, 2.5 µg/µl Collagenase D in culture medium) at 37°C. The resulting MEF and MPF suspensions were separated from undigested tissues using a cell strainer and transferred into culture medium comprising DMEM (Thermo Fisher Scientific, Germany), 10% fetal bovine serum (Bio&SELL, Germany), 1% MEM non-essential amino acids (Sigma-Aldrich, Germany), 1% Cell Culture Guard (AppliChem, Germany) and 2 mM L-Glutamine (Sigma-Aldrich, Germany). Cells were cultured in a humidified atmosphere with 5% CO2 at 37°C and split regularly until growth ceased (passages ∼ 6 to 12). Culture medium was then replaced once every two weeks until cells spontaneously immortalized and resumed proliferation (∼4-12 weeks post growth stop).

### Transfection

siRNAs were diluted in OptiMEM (Thermo Fisher Scientific, Germany) to a final concentration of 100 nM. After adding 6 µl of Lipofectamine RNAiMAX (Thermo Fisher Scientific, Germany), the solution was incubated for 5 min at RT, and then added dropwise onto freshly seeded cells. Cells were incubated with siRNA/lipid complexes for 72 h under standard conditions. The following siRNAs (sense sequence, 5’-3’) were utilized: AllStars Control siRNA (siCtrl, Qiagen, Germany); siRad54l (cca ccu ggu uau aac ucu att, Thermo Fisher Scientific, Germany); siNek1 (siRNA pool containing the four sequences aua gcu aaa cga auc gaaa, ggu gau uug uuu aaa cgaa, gcu cga acu ugc aua ggca, and ggg aua gac cau cag ucaa, Dharmacon, USA). For targeting other Nek kinases, individual siRNAs were employed, all obtained from Thermo Fisher Scientific, Germany: siNek2 (cau cgu cag uua cua uga utt); siNek3 (gau ugg gug uca gaa cua att); siNek4 (gaa gca cgc uuu uaa ugc att); siNek5 (cau gaa gac uau aca gac att); siNek6 (cca ccg acc uga cau ugu att); siNek7 (gga uaa uga gcu gaa cau att); siNek8 (ccu ucg cac cca ucu cug att); siNek9 (cca uuc uua uug uug aga att); siNek10 (cca aug uug uac guu auu att); siNek11 (gga cca aga uga agc gca utt). Plasmids were introduced into cells 24 h after seeding or siRNA transfection using Lipofectamine 2000 (Thermo Fisher Scientific, Germany) following the manufacturer’s instructions. Briefly, 10 µg of plasmid DNA was incubated with 6 µl of Lipofectamine 2000 (Thermo Fisher Scientific, Germany) in OptiMEM at RT for 30 min. Meanwhile, cells were provided with transfection medium comprising DMEM (Thermo Fisher Scientific, Germany), 10% fetal bovine serum (Bio&SELL, Germany), 1% MEM non-essential amino acids (Sigma-Aldrich, Germany) and 2 mM L-Glutamine (Sigma-Aldrich, Germany). The plasmid/lipid complexes were added dropwise to cells and left on for 6 h under standard conditions. Cells were then provided with fresh transfection medium and cultured under standard conditions for 48 h.

### DNA damage induction

10 µM EdU was added to cells 1 h before irradiation and maintained throughout the repair incubation period to allow for the identification of S-phase cells in subsequent analyses. DNA damage, including DNA double-strand breaks (DSBs), was induced by irradiating the cells with a Titan E Isovolt 160 X-ray tube (GE Technologies, USA) at 90 kV using a 1 mm aluminum plate and a dose rate of ∼1 Gy/min. For cells seeded on glass coverslips, a dose correction factor of 3 was considered to correct for secondary radiation effects from the glass.^65^

### γH2ax and Rad51 foci assays

Cells were seeded onto glass coverslips, treated as indicated in the figures and then fixed with 4% paraformaldehyde (Sigma-Aldrich, Germany) for 15 min at RT. After washing with PBS, cells were permeabilized with 0.3% Triton-X100/PBS for 5 min at RT and incubated in 1x Roti-Immuno-Block (Roth, Germany) for 30 min at RT. Primary antibodies were diluted in 1x Roti-Immuno-Block and added to the cells, which were then incubated overnight at 4°C in a humid chamber. Following incubation, cells were washed with 0.1% Tween20/PBS and incubated with secondary antibodies diluted in 1x Roti-Immuno-Block for 1 h at RT in the dark. The following antibodies were used for Immunofluorescence: Chicken anti-GFP (Abcam, 1:400- 1:1000); Goat anti-Rad54 (Santa Cruz Biotechnology, 1:400); Mouse anti-phospho-H2ax (Merck Millipore, 1:1000); Rabbit anti-mCherry (Abcam, 1:400); Rabbit anti-pH3 (Merck Millipore, 1:1000); Rabbit anti-Rad51 (Abcam, 1:10000); Donkey anti-chicken Alexa Fluor488 (Dianova, 1:1000); Donkey anti-mouse Alexa Fluor 488 (Invitrogen, 1:1000); Donkey anti-rabbit Alexa Fluor 488 (Thermo Fisher Scientific, 1:1000); Donkey anti-goat Alexa Fluor 647 (Molecular Probes, 1:200); Donkey anti-mouse DyLight550 (Abcam, 1:1000); Donkey anti-rabbit DyLight550 (Thermo Fisher Scientific, 1:1000). After washing with 0.1% Tween20/PBS, cells were stained for EdU using the Click-iT EdU Imaging Kit (Baseclick, Germany) as instructed by the manufacturer. Finally, cells were washed with PBS, treated with a 0.4 µg/ml DAPI solution (Sigma-Aldrich, Germany) for 5 min at RT to stain DNA and mounted onto glass slides. Slides were examined on an Axiovert 200M microscope with AxioCamMRm (Zeiss, Germany). Metafer4 (MetaSystems) was used to scan the slides at 10x magnification thereby plotting DAPI and EdU signal intensities in a diagram. Based on the resulting distribution in the diagram, G2-phase cells were distinguished from G1-phase cells based on their doubled DNA content and, thus, doubled DAPI signal. pH3 served as an additional marker for G2-phase cells. S-phase cells were differentiated from the G1/G2 population by their high EdU signal. DNA damage repair was evaluated by counting γH2ax or Rad51 foci in 50 EdU-negative, pH3-positive G2 cells at the microscope at 100x magnification. Immunofluorescent images in the figures were captured using an Imager.Z2 microscope with AxioCamMRm (Zeiss, Germany) or an Axioimager M1 microscope with ApoTome.2 (Zeiss, Germany) at 100x magnification and processed with Fiji or ZEN Blue (Zeiss, Germany, version 2.6).

### Chromatid breaks

Cells were incubated with 50 ng/ml Calyculin A (LC Laboratories, USA) for 30 min at standard conditions, then collected at indicated timepoints and centrifuged at 180 g for 10 min at 4°C. Prewarmed 75 mM KCl was added dropwise to the cells, followed by 12-min incubation at 37°C. A few drops of fresh fixative (3 parts methanol:1 part glacial acetic acid) were added before centrifuging at 150 g for 10 min at 4°C. Cells were then resuspended in fixative, the suspension was incubated for 10 min at RT and centrifuged again. The fixation procedure was repeated two more times. Cell suspensions were dropped onto slides and the resulting chromosome spreads were stained using 5% Giemsa (Sigma-Aldrich, Germany) for 20 min at RT, followed by several washes with distilled water. After drying, chromosomes were scanned and imaged using an Imager.Z2 microscope with AxioCamMRm (Zeiss, Germany) equipped with the Metafer4 software (MetaSystems, Germany) at 10x and 100x magnification. Images were processed with Fiji and chromatid breaks were counted in at least 40 spreads per condition using the Cell Counter Plugin.

### SCE measurements

24 h after siRNA transfection, cells were incubated with 10 µM BrdU (BD Biosciences, USA) for 36 h under standard conditions and then irradiated as described above. 100 nM Chk1-inhibitor UNC-01 (Merck, Germany) and 0.2 µg/ml colcemid (Sigma-Aldrich, Germany) were added 3 h before harvesting. Suspensions were centrifuged, treated and dropped onto slides as described for the chromatid break assay. The resulting chromosomal spreads were stained with 5 µg/ml Bisbenzimide for 1 h in the dark, washed and dried at 37°C. After covering the spreads with IR-buffer (200 mM Na2HPO4, 4 mM citric acid), they were irradiated with ultraviolet light C at 9 Joules/cm², washed and incubated in 2x saline-sodium citrate buffer (300 mM NaCl, 30 mM sodium citrate, pH 7) for 30 min at 55°C. Chromosomes were stained with Giemsa and at least 40 spreads per condition were analyzed as described above.

### Quantitative real-time PCR (qPCR)

RNA was isolated from cells according to the protocol of the MasterPure Complete DNA and RNA Purification kit (Lucigen, USA). 1 µg of RNA was reverse transcribed into cDNA using the RevertAid First Strand cDNA Synthesis kit as instructed by the manufacturer (ThermoFisher Scientific, Germany). qPCR was prepared with 1:3 diluted cDNA, FastStart Universal SYBR Green Master Mix (Merck, Germany) and 0.2 µM of the respective primers before amplification in a StepOnePlus light cycler (ThermoFisher Scientific, Germany) under the following conditions: 95°C for 10 min, 40 cycles at 95°C for 15 sec and 58°C for 60 sec. A melting curve was staged right after each amplification process to confirm the specificity of all primer pairs. qPCR data were collected from technical duplicates. Fold changes in expression levels were calculated based on the resulting cycle thresholds and the 2^-ΔΔCt^ method in which *Gapdh* served as reference gene. The following primers (forward, reverse; 5’-3’) were used for amplification: Gapdh (cag caa gga cac tga gca aga, tat ggg ggt ctg gga tgg aaa); Nek1 (ttg gac cac agc ctc tccca, cgg gaa ttt gat ggg cct gtt); Nek2 (aag ctg ggg gac ttt ggact, ggg agg cat tag tgc aca cag); Nek3 (aga gca gcc aga gga aat cca, cag acc cac ctc tgt cat cctc); Nek4 (cac tac cag cca cgc tct tct, ctt gtt tgc cag tgt ggc ctt); Nek5 (ggc tag gat gga gca tcc caat, aca cag gat ctg gtc ttc gct); Nek6 (gcg ggt gac ctc tca cagat, ttg gcg ggc ttg atg tctcg); Nek7 (cac aag gaa tgc aag ggccg, gct acc ggc act cca tccaa); Nek8 (cat caa gca tgt ggc ctgcg, ttc tac aat ggt ggg ctg gct); Nek9 (gaa tat gga cgg ctg ggt ttgg, agg tcc cat cac agc cac att); Nek10 (tgc aag tgg agc cca caaga, cgg cag tga gtc ttt tgg agag); Nek11 (tcc ttc att gag acc gtc ggc, ggt gaa gct ttt tct gca cggc).

### SDS-PAGE and immunoblotting

Cells were harvested in RIPA buffer (50 mM Tris pH 8, 150 mM NaCl, 0.1% SDS, 0.5% sodium deoxycholate, 1% Triton-X100, 1x cOmplete protease inhibitor cocktail, 1 mM Na3VO4, 2 mM NaF). The resulting suspension was incubated on ice for 1 h. The protein-containing fraction was separated from cellular debris by centrifugation for 15 min at 4°C and 18,000 g. 40 µg of protein was supplemented with 6x reducing loading buffer (350 mM Tris pH 6.8, 10% SDS, 50% Glycerol, 600 mM DTT, 0.05% Bromophenol blue) and denatured by heating for 5-10 min at 95°C. Protein samples were loaded onto an SDS gel (5% stacking, 8% separation), separated at 90 mA for at least 1.5 h and finally blotted onto a nitrocellulose membrane with a current of 310 mA for 3 h. The membrane was cut at desired protein sizes and blocked with 5% skim milk (Roth, Germany) for 1 h at RT before incubation with primary antibodies diluted in 5% BSA (PAN-Biotech, Germany) at 4°C overnight. After washing with TBS-T (20 mM Tris pH 7.6, 137 mM NaCl, 0.1% Tween20), the membrane was incubated with secondary antibodies diluted in 5% skim milk for 1 h at RT. The following antibodies were used for staining: Mouse anti-Gapdh (Santa Cruz Biotechnology, 1:200000); Mouse anti-Nek1 (Santa Cruz Biotechnology, 1:500); Mouse-anti GFP (Roche, 1:500); Mouse anti-Rad54 (Santa Cruz Biotechnology, 1:200); Rabbit anti-Rad51 (Abcam, 1:2500); Donkey anti-mouse-HRP (Dianova, 1:1000); Donkey anti-rabbit-HRP (Dianova, 1:1000). Protein bands were visualized with the Dual-mode Imaging System Fusion FX (Vilber Lourmat, Germany) after incubation of the membrane with the WesternBright Quantum blotting substrate (Advansta, USA).

### Statistics

Unless otherwise indicated, data were collected from three independent experiments and are presented in figures as mean values with error bars representing the standard error of the mean (SEM). For animal studies, each mouse was considered as one individual experiment. Diagrams and statistical analyses were generated using OriginPro2024 (OriginLab, USA). One-way ANOVA followed by Bonferroni correction for multiple comparisons was performed to determine statistical significance. A p-value of less than 0.05 was considered statistically significant.

## Acknowledgements

We thank Ratna Cordoni, Cornelia Schmitt and Bettina Basso for excellent technical support, Jasmin Messerer and Carmen Schwarzer for help with the mouse experiments, and Bodo Laube for support in the design of the mouse studies. This work was funded by the Federal Ministry of Research, Technology and Space (02NUK034A and 02NUK054C), the Deutsche Forschungsgemeinschaft (DFG, German Research Foundation) – Project-ID 393547839 – SFB 1361, and the DFG GRK1657.

