## Supplemental figures for "Nek family members regulate Rad54 during homologous recombination in developing mice"

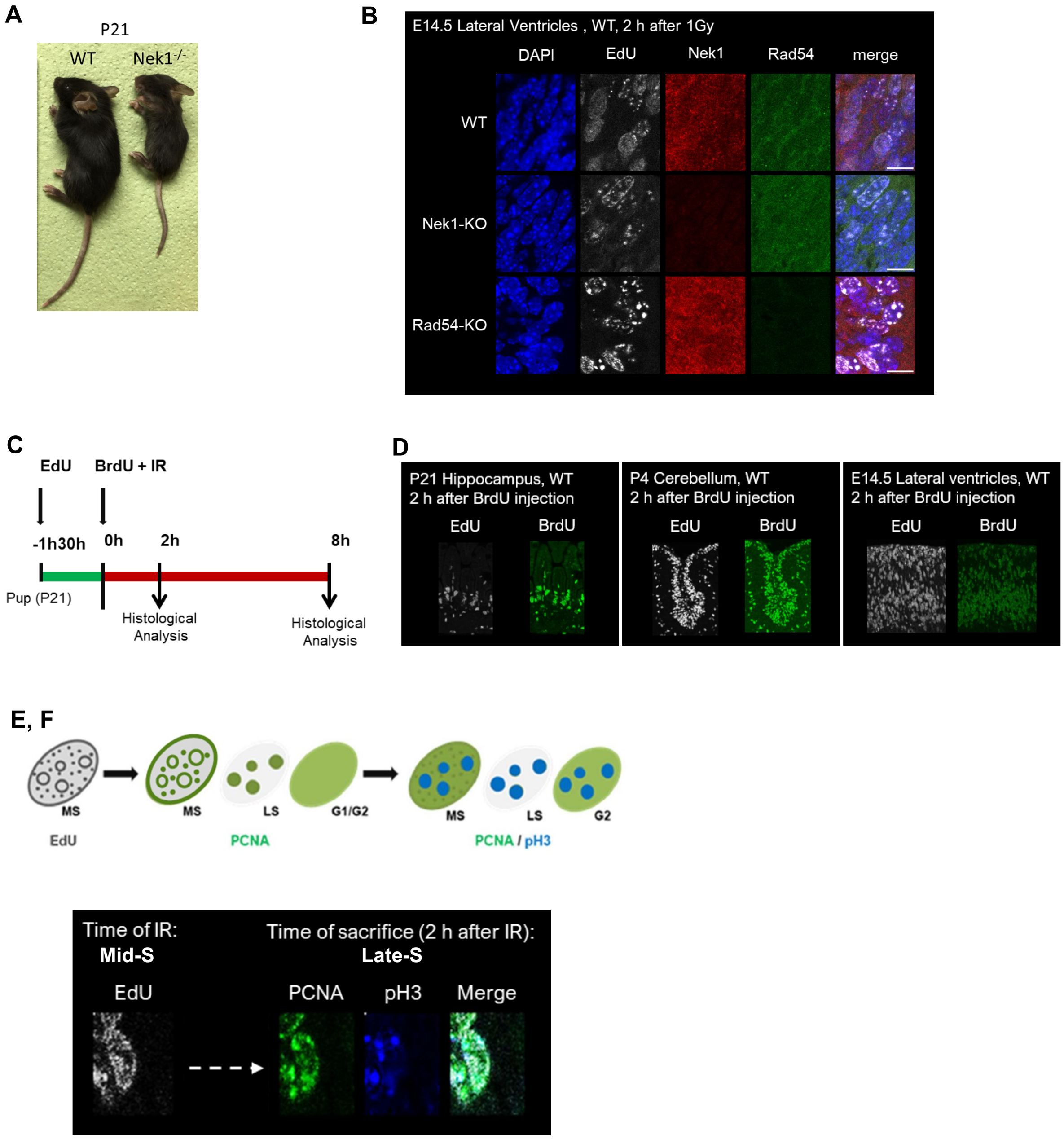


*Fig. S1: In vivo experimental scheme and staining procedure.*

A: Representative image of a male 21-day-old Nek1-KO mouse (Nek1$^{-/-}$, right) compared to a WT mouse of the same sex and age (left). As previously reported, Nek1-KO mice show a marked growth retardation, which can be observed from day 14 after birth and manifests itself clearly in the following 7 days.

B: Representative immunofluorescence images of brain sections demonstrating the presence of Nek1 and Rad54 in irradiated WT, Nek1-KO and Rad54-KO mouse embryos. Mice were treated as described in C and tissues were stained for DNA (DAPI), EdU, Rad54 and Nek1 using specific kits and antibodies.

C: Schematic diagram of the animal experiment. Mice were treated with 3.8 g/ml of the thymidine analogue EdU via intraperitoneal injection to label replicating cells in S phase. 1.5 h later, mice were similarly treated with 7.6 g/ml of the thymidine analogue, BrdU, which preferentially incorporates into

DNA over EdU. This “washout” of EdU allows the detection of cells that were located in mid-S phase at the time of irradiation. Mice were irradiated with 1 Gy of X-rays immediately after BrdU injection and sacrificed 2 h or 8 h later. Brains or small intestines were removed, fixed in formaldehyde for 24 h, embedded in paraffin and subjected to histological and immunofluorescence stainings.

D: Representative immunofluorescence images of brain sections from postnatal (left, P21), neonatal (middle, P4) and embryonic (right, E14.5) WT mice demonstrating the preferential incorporation of BrdU over EdU. Mice were treated as described in C. Sections with a thickness of 4 microns were prepared and stained for EdU and BrdU.

E: Schematic diagram of the staining procedure to identify cells that were irradiated in mid-S phase and to determine their cell cycle phase at the time of fixation. Mid-S phase irradiated cells were identified by their specific EdU staining pattern. PCNA was then stained to distinguish S phase cells at the time of sacrifice. As a S-phase marker, PCNA cannot be used to discriminate between G1 and G2 phase cells. Phospho-Histone 3 was therefore stained to specifically label G2 phase cells. MS, mid-S; LS, late-S

F: Representative immunofluorescence images of the staining procedure described in E.

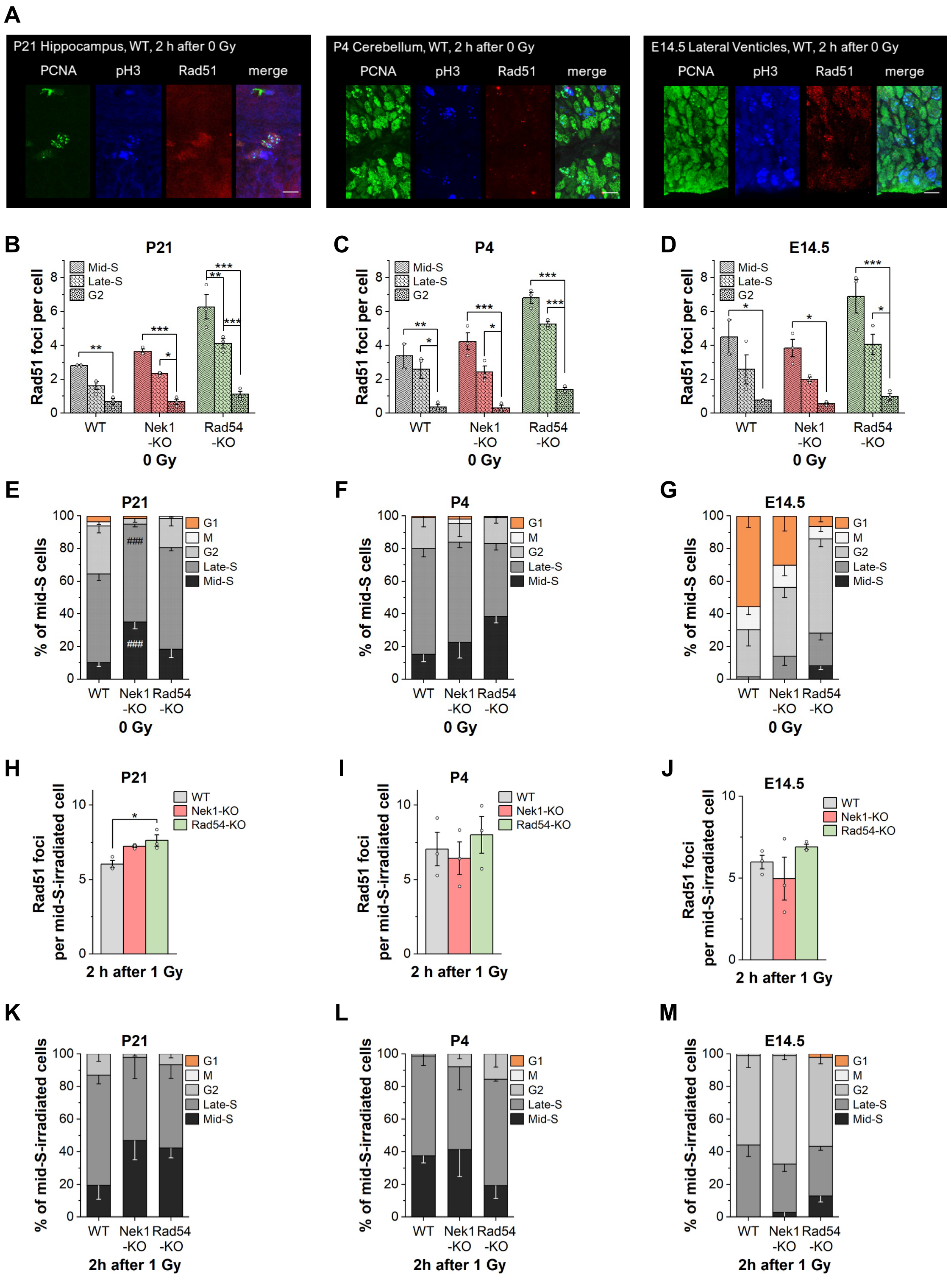


*Fig. S2: Rad51 foci assay in brain tissues - unirradiated control and 2 h after 1 Gy samples.*

A: Representative immunofluorescence images of brain sections from unirradiated postnatal (A, P21), neonatal (B, P4) and embryonic (C, E14.5) WT mice. Mice were treated with EdU by intraperitoneal (i.p.) injection for 1.5 h and sacrificed 2 h or 8 h after BrdU injection. Brains were isolated, fixed in formaldehyde for 24 h and embedded in paraffin. Sections with a thickness of 4 microns were prepared and stained for cell cycle markers EdU, PCNA, phospho-Histone 3 (pH3) and the HR marker Rad51 using specific kits or antibodies, respectively.

B, C, D: Rad51 foci assay in brain tissues of unirradiated postnatal (A), neonatal (B) and embryonic (C) mice which were treated as described in A. Mid-S phase, late-S phase and G2 phase cells in the hippocampus (A), cerebellum (B) or lateral ventricles (C) were detected according to their PCNA and pH3 staining pattern, and then analyzed for their levels of spontaneously arising Rad51 foci.

E, F, G: Cell cycle analysis in brain tissues of unirradiated postnatal (D), neonatal (E) and embryonic (F) mice. Mice were treated as described for panel A, B, C. The cycle phase of cells was identified based on the cellular staining patterns for PCNA (S phase marker) and pH3 (G2 phase marker).

H, I, J: Rad51 foci assay in brain tissues of irradiated postnatal (G), neonatal (H) and embryonic (I) mice. Mice were treated with 3.8 g/ml EdU by intraperitoneal (i.p.) injection, exposed to 1 Gy and sacrificed 2 h later followed by a treatment corresponding to panels A, B, C. Rad51 foci were quantified in all cells that, based on their EdU pattern, were irradiated in mid-S phase. When comparing Rad51 foci data of this induction time point with those from the repair time point (see Figs. 1 D-F), it may appear that the majority of HR events were efficiently repaired in all genotypes 8 h after IR. However, it should be noted that foci levels comprise both, spontaneous (ca. 0.5 to 6 depending on developmental stage, cell cycle phase and genotype; see B-D) and IR-induced Rad51 foci. The significant decrease in the total number of foci is therefore also related to the transition of cells into a cell cycle phase with fewer spontaneous DSBs.

K, L, M: Cell cycle analysis in brain tissues of irradiated postnatal (J), neonatal (K) and embryonic (L) mice. Mice were treated as described for panel G, H, I. The cycle phase of irradiated mid-S phase cells was identified based on the cellular staining patterns for PCNA (S phase marker) and pH3 (G2 phase marker).

Data are shown as mean values of at least 2 animals ± SEM (n≥2). White circles indicate results from individual mice. $^{*,\#}p < 0.05$; $^{**,\#\#}p < 0.01$; $^{***,\#\#\#}p < 0.001$ (ANOVA).

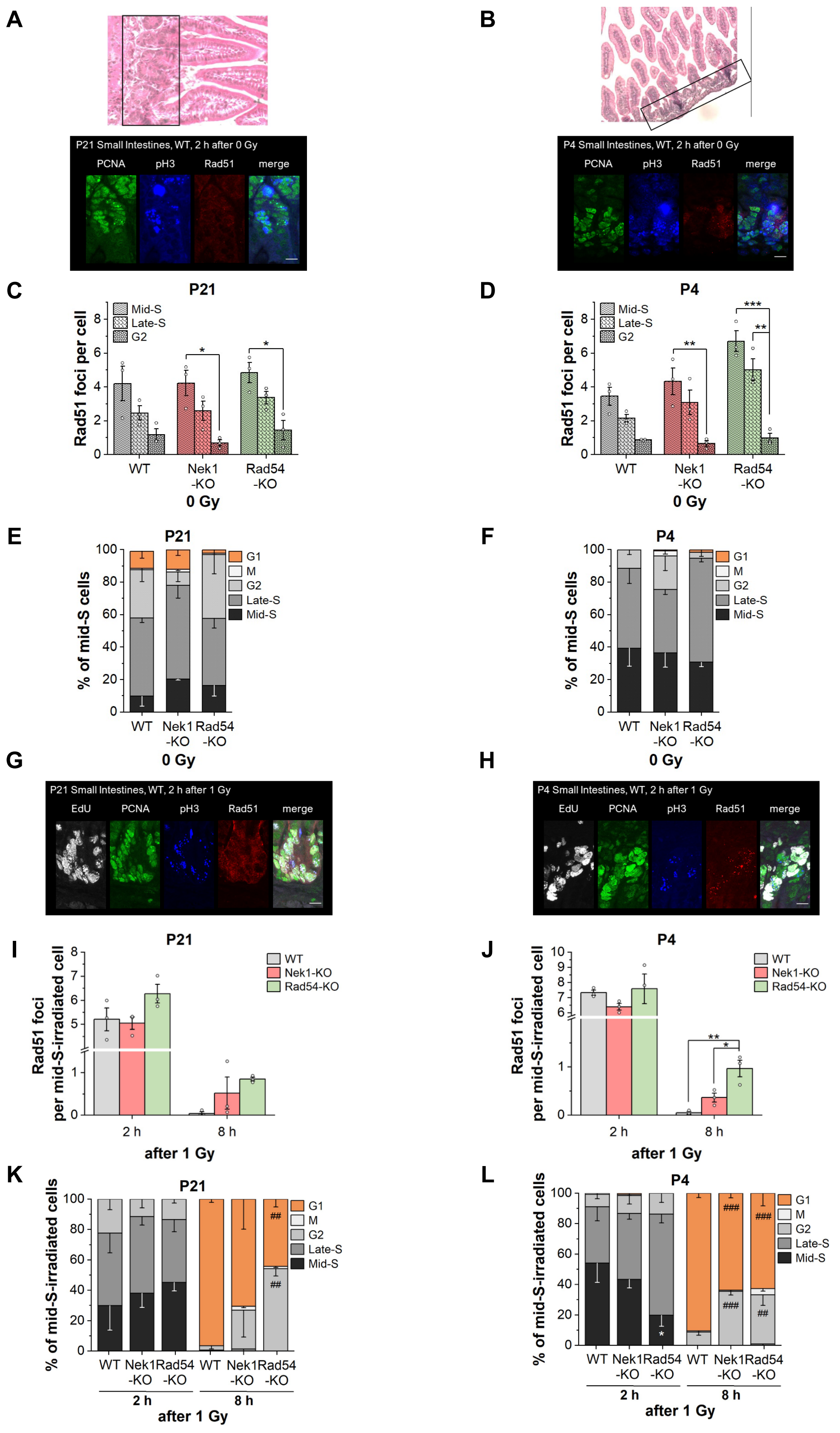


*Fig. S3: The involvement of Nek1 in HR changes during small intestine development in vivo.*

A, B: Representative histological (top) and immunofluorescence (bottom) images of small intestine sections from unirradiated postnatal (A, P21) and neonatal (B, P4) WT mice. Mice were treated with EdU by intraperitoneal (i.p.) injection for 1.5 h and sacrificed 2 h or 8 h after BrdU injection. Small intestines were isolated, fixed in formaldehyde for 24 h and embedded in paraffin. Sections with a thickness of 4 microns were prepared and stained for cellular structures using hematoxylin and eosin (top) or for cell cycle markers EdU, PCNA, phospho-Histone 3 (pH3) and the HR marker Rad51 using specific kits or antibodies (bottom). Black boxes indicate area within the crypts with proliferative cell populations. Scale bars represent 10 µm.
C, D: Rad51 foci assay in the small intestines of unirradiated postnatal (C) and neonatal (D) mice which were treated as described in A, B. Mid-S phase, late-S phase and G2 phase cells in the crypts of the small intestines were detected according to their PCNA and pH3 staining pattern, and then analyzed for Rad51 foci levels.
E, F: Cell cycle analysis in the small intestines of unirradiated postnatal (E) and neonatal (F) mice which were treated as described in A, B. The cycle phase of cells was identified based on the cellular staining patterns for PCNA (S phase marker) and pH3 (G2 phase marker).
G, H: Representative immunofluorescence images of small intestine sections from postnatal (G) and neonatal (H) WT mice. Mice were treated with EdU by intraperitoneal (i.p.) injection, exposed to 1 Gy of X-rays 1.5 h later and sacrificed 2 h or 8 h after IR. Small intestines were isolated, fixed in formaldehyde for 24 h and embedded in paraffin. Sections with a thickness of 4 microns were prepared and stained for cell cycle markers EdU, PCNA, phospho-Histone 3 (pH3) and the HR marker Rad51 using specific kits or antibodies, respectively.
I: Rad51 foci assay in small intestine tissue of postnatal mice. Mice were treated as described for panel G, H. Rad51 foci were quantified 2 and 8 h after IR and in all cells that were in the areas described in A, B and irradiated in mid-S phase according to their EdU staining pattern. When comparing Rad51 foci data of the induction time point (2 h after IR) with those from the repair time point (8 h after IR), it may appear that most HR events were efficiently repaired in all genotypes. However, it should be noted that foci levels comprise both spontaneous (ca. 0.5 to 6 depending on developmental stage, cell cycle phase and genotype; see C, D) and IR-induced Rad51 foci. The significant decrease in the total number of foci is therefore also related to the transition of cells into a cell cycle phase with fewer spontaneous DSBs.
J: Rad51 foci assay in small intestine tissue of neonatal mice. Mice were treated as described for panel G, H. Rad51 foci were quantified 2 and 8 h after IR with 1 Gy and in all cells that were in the areas described in A, B and irradiated in mid-S phase according to their EdU staining pattern.
K, L: Cell cycle analysis in small intestine tissue of postnatal (K) and neonatal (L) mice. Mice were treated as described for panel G, H. Cycling of irradiated mid-S phase cells was evaluated based on their cellular staining patterns for PCNA (S-phase marker) and pH3 (G2-phase marker).
Data are shown as mean values of at least 2 animals ± SEM (n≥2). White circles indicate results from individual mice. $^{*,\#}p < 0.05$; $^{**,\#\#}p < 0.01$; $^{***,\#\#\#}p < 0.001$ (ANOVA).

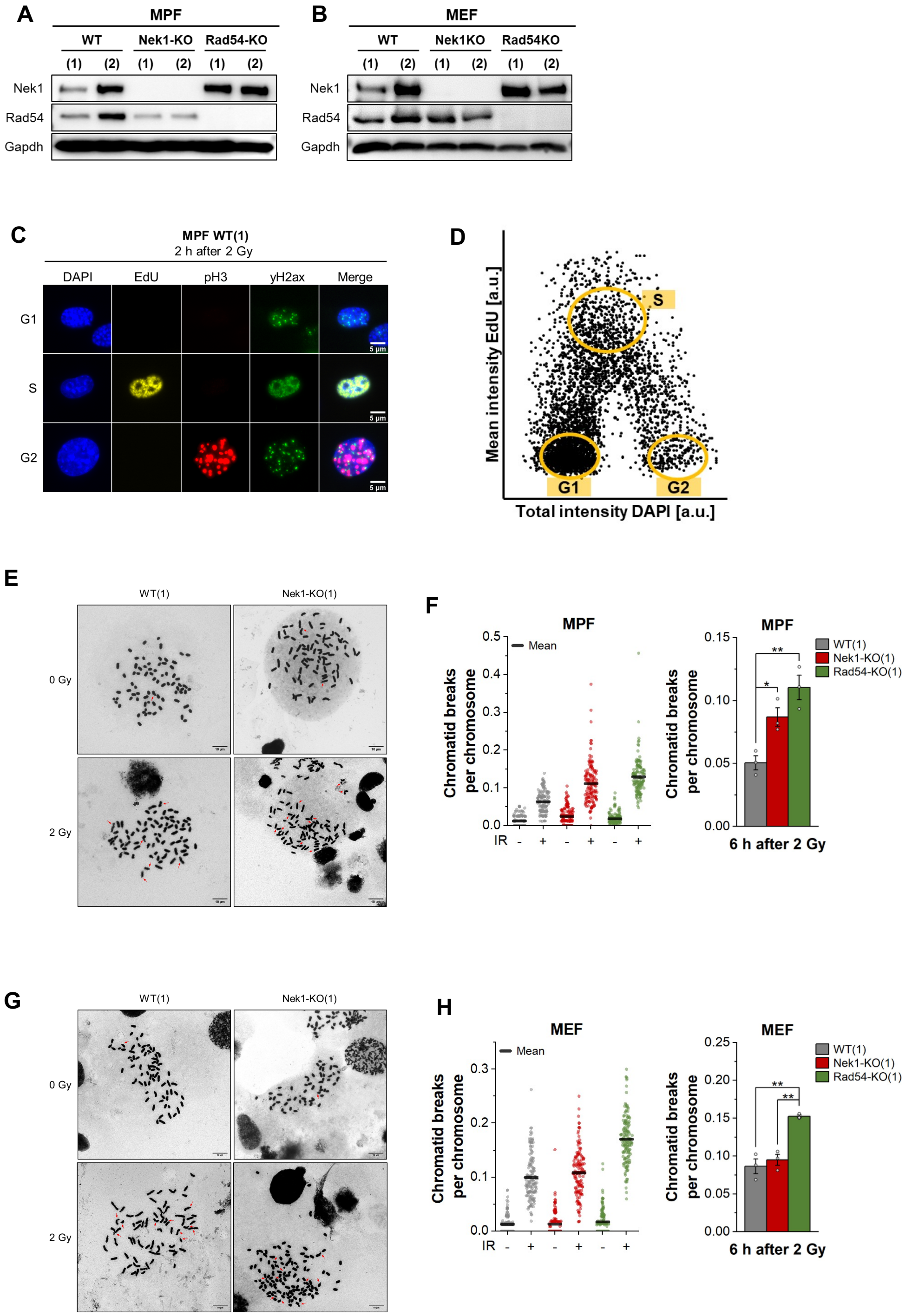


*Fig. S4: In vitro staining and analysis procedure in MPF and MEF samples and chromosomal analysis.*

A, B: Representative Western Blots showing the protein levels of Nek1 and Rad54 in MPFs (A) and MEFs (B) isolated from two individual postnatal (MPFs) or embryonic (MEFs) WT, Nek1-KO or Rad54-KO mice (1, 2). Gapdh served as loading control.

C: Representative fluorescence images of the γH2ax- foci assay. Fibroblasts were treated with 10 µM EdU for 1 h and then irradiated with 2 Gy, fixed 2 h after IR, and stained for DNA (DAPI), EdU as S phase marker, phospho-Histone 3 (pH3) as G2 phase marker, and γH2ax labeling double-strand breaks. To allow the

precise analysis of DNA repair by HR, γH2ax foci were quantified in EdU-negative, pH3-positive cells as they were in G2 phase at the time of irradiation.
D: The cell cycle distribution was recorded using the Metafer4 software (Metasystems). The intensities (a.u.) for EdU and DAPI were plotted into a histogram to identify EdU-positive S phase cells and to distinguish EdU-negative G1 and G2 phase cells by their DNA content (DAPI low = G1 phase, DAPI high = G2 phase cells). G2 phase cells were additionally labeled with pH3.
E: Representative chromosome spreads of unirradiated or irradiated WT and Nek1-KO MPFs.
F: Chromatid breaks in WT, Nek1-KO and Rad54-KO MPFs. Fibroblasts were fixed 6 h after IR with 2 Gy or left unirradiated and chromosome spreads were obtained from G2-phase cells.
G: Representative chromosome spreads of unirradiated or irradiated WT and Nek1-KO MEFs.
H: Chromatid breaks in WT, Nek1-KO and Rad54-KO MEFs. Fibroblasts were fixed 6 h after IR with 2 Gy or left unirradiated and chromosome spreads were obtained from G2-phase cells.
Data are shown as individual results for the analysis of at least 120 cells (left) per condition or as mean values ± SEM (right) (n=3). White circles indicate results from individual experiments. Spontaneous chromatid breaks were subtracted from the irradiated samples. *$p < 0.05$; **$p < 0.01$; ***$p < 0.001$ (ANOVA). MPF, murine postnatal fibroblasts; MEF, murine embryonic fibroblasts.

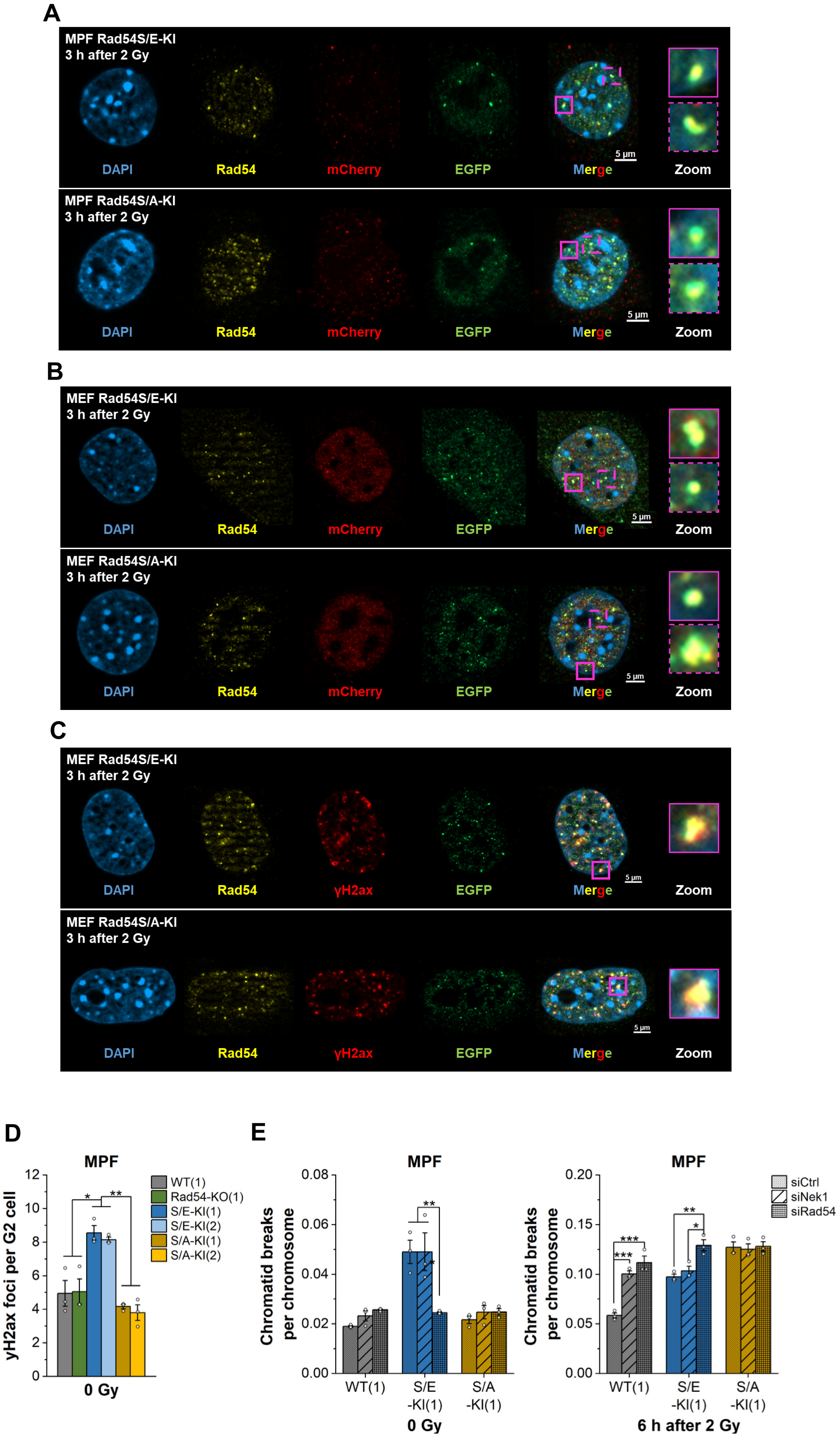


*Fig. S5: Functional validation KI MEFs (colocalization), spontaneous foci and chromatid breaks in MPFs.*

A: Representative images of expression studies in Rad54S/E-KI (top) and S/A-KI (bottom) MPFs. Fibroblasts were irradiated with 2 Gy, fixed 3 h thereafter and stained for DNA (DAPI, blue), Rad54 (yellow), mCherry (red), and EGFP (green). Examples of Rad54-GFP colocalizations are depicted in a magnified version (Zoom). These images confirm that Rad54-KI fibroblasts do express the GFP-tagged Rad54S/E or S/A variant instead of the mCherry-tagged Rad54 WT protein. Images were taken at a 1000x magnification using an Axioimager M1 microscope equipped with an ApoTome.2.

B, C: Representative images of expression (B) and colocalization (C) studies in Rad54S/E-KI (top) and S/A-KI (bottom) MEFs. Fibroblasts were irradiated with 2 Gy, fixed 3 h thereafter and stained for DNA (DAPI, blue), Rad54 (yellow), mCherry (red), and EGFP (green) in B or for DNA (DAPI, blue), Rad54 (yellow), γH2ax (red), and EGFP (green) in C. Examples of Rad54-GFP (B) and Rad54-GFP-γH2ax (C) colocalizations are depicted in a magnified version (Zoom). Images were taken at a 1000x magnification using an Axioimager M1 microscope equipped with an ApoTome.2.

D: γH2ax foci assay in unirradiated MPFs isolated from Rad54S/E-KI or S/A-KI (1, 2), WT (1) or Rad54-KO (1) postnatal mice. Fibroblasts were treated with 10 µM EdU for 1 h, fixed 2 h thereafter and stained for γH2ax, phospho-Histone 3 (pH3) and EdU.

E: Chromatid breaks in Nek1- and Rad54-depleted WT and Rad54-KI MPFs. 72 h after transfection with 100 nM unspecific control, Nek1- or Rad54-specific siRNA, fibroblasts were irradiated with 2 Gy, or left unirradiated, and fixed 6 h thereafter and chromosome spreads were obtained from G2-phase cells.

Data are shown as mean values ± SEM (n=3). White circles indicate results from individual experiments. Spontaneous chromatid breaks were subtracted from the irradiated samples. *$p < 0.05$; **$p < 0.01$; ***$p < 0.001$ (ANOVA). MPF, murine postnatal fibroblasts; MEF, murine embryonic fibroblasts.

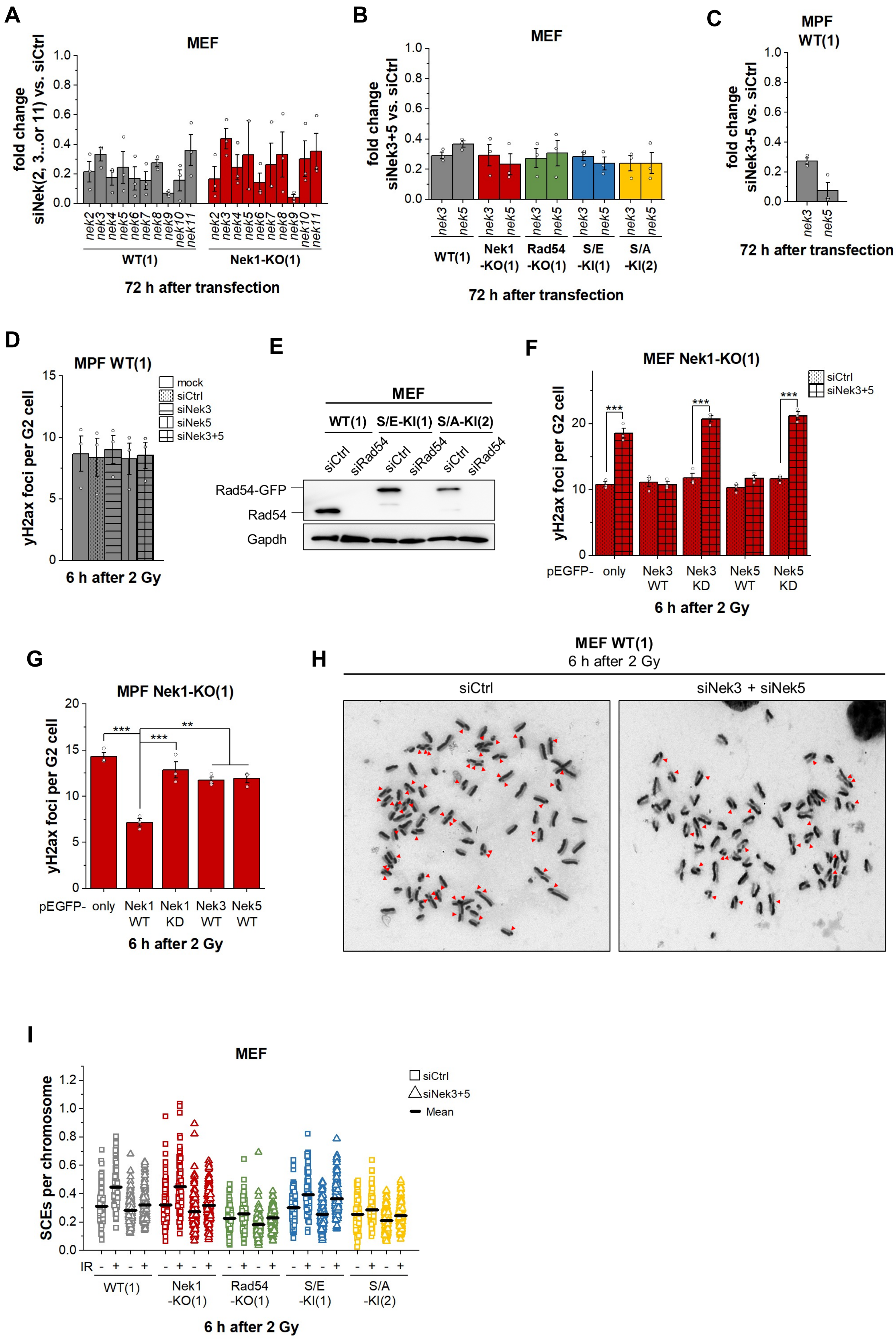


*Fig. S6: Knockdown validation of Nek-specific siRNAs with qPCR, γH2ax foci assay in Nek3/5-depleted MPFs, rescue assays in Nek1-KO MEFs and Nek1-KO MPFs, SCEs in Nek3/5-depleted MEFs.*

A: Gene expression analysis in WT and Nek1-KO MEFs following siRNA transfection. Fibroblasts were treated with 100 nM unspecific control or Nek-specific siRNA for 72 h at standard conditions. Fold changes in expression were calculated with the $2^{-\Delta\Delta Ct}$ method using *Gapdh* as a reference gene and normalizing the expression levels of Neks in fibroblasts treated with Nek-specific siRNAs to their respective level in fibroblasts treated with siCtrl.
B, C: Gene expression levels of Nek3 and Nek5 in WT, Nek1-KO, Rad54-KO and Rad54-KI MEFs (B) as well as in WT MPFs (C) following double transfection with 100 nM Nek3- and Nek5-specific siRNAs for 72 h at standard conditions. Values are depicted as fold change to samples treated with siCtrl.
D: γH2ax foci assay in WT MPFs. 72 h after Nek3- and Nek5-siRNA transfection as described for panel C, fibroblasts were treated with 10 µM EdU for 1 h, irradiated with 2 Gy, fixed 6 h thereafter and stained for γH2ax, phospho-Histone 3 (pH3) and EdU.
E: Representative Western Blot showing the protein levels of Rad54 in Rad54-KI MEFs following transfection with 100 nM unspecific control or Rad54-specific siRNA for 72 h at standard conditions. Gapdh served as loading control.
F: Quantification of rescue assays in Nek1-KO MEF as shown in Fig. 4E. Fibroblasts were treated with 100 nM unspecific control or Nek3- and Nek5-specific siRNAs for 24 h at standard conditions followed by transfection with 10 µg of plasmids coding for only GFP, the wildtype forms of Nek3 or Nek5 (WT) or kinase-dead variants of Nek3 or Nek5 (KD). 48 h after plasmid transfection, fibroblasts were treated with 10 µM EdU for 1 h and then irradiated with 2 Gy, fixed 6 h thereafter and stained for γH2ax, phospho-Histone 3 (pH3), GFP and EdU.
G: γH2ax foci assay in Nek1-KO MPFs. Fibroblasts were transfected with 10 µg of plasmids coding for only GFP, the wildtype form of Nek1, Nek3 or Nek5 (WT) or a kinase-dead variant of Nek1 (KD). 48 h after plasmid transfection, fibroblasts were treated with 10 µM EdU for 1 h and then irradiated with 2 Gy, fixed 6 h thereafter and stained for γH2ax, phospho-Histone 3 (pH3), GFP and EdU.
H: Representative images of SCEs in chromosome spreads from siCtrl- or siNek3- and siNek5-treated WT MEFs 6 h after IR. Fibroblasts were irradiated with 2 Gy, or left unirradiated, and fixed 6 h thereafter and chromosome spreads were obtained from mitotic cells.
I: Quantification of SCEs as shown in panel H.
Data are shown as mean values ± SEM (n=3). White circles indicate results from individual experiments which in case of the γH2ax assay corresponds to the quantification of 50 EdU-negative, pH3-positive G2 phase cells per experiment or in case of SCE measurements corresponds to the quantification of at least 40 chromosome spreads per experiment. Spontaneous foci were subtracted from the irradiated samples. *$p < 0.05$; **$p < 0.01$; ***$p < 0.001$ (ANOVA). MPF, murine postnatal fibroblasts; MEF, murine embryonic fibroblasts.
